# Developmental regulator-mediated transcriptional reprogramming reveals novel regulators of somatic embryogenesis in sorghum

**DOI:** 10.64898/2026.11.07.764635

**Authors:** Pallavi Dhiman, Leonidas D’Agostino, Vaishnavi Deshmukh, Luis Herrera-Estrella, Madhusudhan Janga, Yinping Jiao, Gunvant B Patil

**Affiliations:** Institute of Genomics for Crop Abiotic Stress Tolerance, Department of Plant and Soil Science, Texas Tech University, Lubbock, TX, 79409, USA

**Keywords:** sorghum, regeneration, developmental regulators, gene-editing, transcriptomics

## Abstract

Recalcitrance to genetic transformation remains a fundamental bottleneck in crop improvement, driven by genotype-specific barriers that include entrenched defense transcriptional states, limited embryogenic competence, and inadequate hormonal reprogramming potential. Here, we present a multi-layered transcriptomic analysis of somatic embryogenesis (SE) and regeneration in two sorghum genotypes (forage Tx2910 and grain Tx2783), two explant types (immature embryos and leaf bases), and three developmental regulator (DR) treatments (GRF-GIF, WUS, and BBM+WUS). This design lets us separate genotype, explant, and DR-specific regulatory programs. Using in-depth transcriptomics analysis (bulk RNAseq and single nucleus transcriptomics), we identified the molecular logic by which DRs override genotype-specific barriers and redirect somatic cells toward embryogenic and regeneration-competent states. Compared to Tx2910, the transformation-recalcitrant sorghum line Tx2783 shows a stronger defense response to both *Agrobacterium* infection plus culture stress and depends more heavily on DR-driven transcriptional reprogramming to exit this defense-locked state. WUS acts as an early, broad suppressor of defense and stress signaling while activating fate-specification programs, whereas GRF-GIF drives a delayed but pronounced transcriptional shift toward cell proliferation and morphogenesis. Integrative gene network analyses further identified a prioritized set of sorghum-specific transcription factors with potential to increase regeneration efficiency. Among the top candidates, we identified that *SbEREB167* and *SbEREB174* (AP2/EREBP family) efficiently induce SE and regeneration in sorghum. Collectively, this study establishes the transcriptional architecture underlying DR-mediated transformation across sorghum types and identifies novel regulators that hold potential to improve plant transformation across recalcitrant cereal species.

## Introduction

Sorghum is a resilient and resource-efficient crop with diverse uses, including food, fuel, fodder, and forage. The rich genetic resources in sorghum provide great potential for improvement, and biotechnological advances have potential to accelerate this trait improvement process (Park *et al*., 2025; Scully *et al*., 2018). While traditional genetics have identified many genes linked to traits such as seed nutrition, stress tolerance, and disease resistance, few have been functionally characterized (Lin *et al*., 2022; Varoquaux *et al*., 2019; Wang *et al*., 2026). High-throughput genotyping, mutant populations, and functional genomics have helped bridge this gap; however, the overall process of trait discovery and deployment in sorghum remains slow. Genetic transformation is an essential tool for both functional genomics and crop improvement, enabling precise manipulation of genes to study their roles and enhance agronomic traits. Sorghum, like many other monocots, has historically been considered recalcitrant to *Agrobacterium*-mediated transformation (Nigam *et al*., 2025).Over the years, plant transformation has advanced from particle bombardment to optimized *Agrobacterium*-mediated methods, improved tissue culture protocols, and the use of DR genes. Recent studies leveraging these advances have led to notable enhancements in sorghum transformation efficiency, paving the way for more effective genetic engineering and trait improvement in this globally important crop (Aregawi *et al*., 2022; Fontanet-Manzaneque *et al*., 2024; Park *et al*., 2025; Wang *et al*., 2023b).

The synergistic expression of DR genes such as *Wuschel2* (*WUS2*), *BabyBoom* (*BBM*), and *GROWTH-REGULATING FACTOR 4 (GRF4)* and its cofactor *GRF-INTERACTING FACTOR 1 (GIF1)* has been demonstrated to enable efficient, genotype-independent transformation in several monocots, including sorghum and recalcitrant maize inbred lines, and has outperformed conventional methods(Debernardi *et al*., 2020; Lowe *et al*., 2016; Nelson-Vasilchik *et al*., 2018). Additionally, these DR-utilizing methods can be effectively implemented in alternative explant sources, such as leaf segments, reducing reliance on immature embryos and shortening the overall tissue culture timeline (Azanu *et al*., 2025; Butler *et al*., 2025; Wang *et al*., 2023b). Additionally, strategies have been developed to mitigate the pleiotropic developmental and reproductive defects caused by constitutive expression of these stem cell developmental regulators, such as Cre/loxP-mediated excision under heat shock or desiccation-inducible promoters. This advance enables the removal of DR genes post-transformation, yielding normal, fertile plants while simultaneously accelerating tissue culture timelines and improving transformation efficiency (Mookkan *et al*., 2017).

Furthermore, advances in sorghum transformation have been largely driven using DR genes combined with optimized vector systems. The ternary vector system, incorporating the pVIR accessory plasmid, enabled improved transformation frequencies in the sorghum reference line Tx430 (6% to 29%) and was successfully adapted to recalcitrant African varieties, facilitating the first stable CRISPR/Cas9-mediated gene knockouts in Tx430 (Che *et al*., 2018; Che *et al*., 2022). Subsequent studies showed that *ZmWUS2* can induce rapid SE formation, enabling the production of high-quality, morphogenic gene-free, and selectable marker-free plants, highlighting the genotype-independent potential of DR-mediated transformation. Beyond Tx430, DRs combined with the ternary vector system markedly improved transformation efficiencies in previously non-transformable sorghum genotypes such as Tx623 and Tx2752 (Che *et al*., 2022). Building on maize leaf transformation strategies, Wang et al. (2023 b) used a maize-optimized WUS2/BBM construct with novel promoter combinations to generate embryogenic callus and plantlets from sorghum leaf explants (Wang *et al*., 2023b).

Further refinements, such as the GRF/GIF ternary vector system pVS1_VIR2, achieved high-efficiency transformation in Tx430 without pleiotropic effects or the need for excision (Li *et al*., 2024). Optimizations in *Agrobacterium* strains, selection regimes, and binary vector design further enhanced transformation efficiencies (Fontanet-Manzaneque *et al*., 2024). Innovative approaches, including viral-mediated, altruistic delivery of DRs, enabled highly efficient leaf-based transformation in Sorghum (Butler *et al*., 2025). Further, scalable seedling leaf whorl transformation mitigated phenolic accumulation and overcame genotype dependence across 13 genetically diverse sorghum genotypes (Azanu *et al*., 2025). Collectively, these studies underscore the transformative impact of DR genes, enabling broader genotype applicability and paving the way for efficient functional genomics and genetic improvement in sorghum.

An underexplored question in this field is whether genotype-dependent recalcitrance reflects a simple quantitative deficit in regenerative capacity or a qualitative transcriptional state dominated by persistent *Agrobacterium*-induced defense signaling and stress-response programs that must be actively overridden before an embryogenic identity can be established.

Distinguishing between these possibilities has direct practical consequences (1) a quantitative deficit could in principle be addressed simply by increasing DR dosage, whereas (2) a distinct, competing transcriptional state would instead require regulators capable of actively suppressing defense/stress programs while simultaneously activating fate-specification and pluripotency programs. In this study, we test the latter hypothesis by comparing the transcriptional trajectories of forage genotype (Tx2910) and grain genotype (Tx2783) under three DR treatments (*ZmBBM*, *ZmWUS2*, and GRF-GIF) and two explant types. By leveraging the ability of known DR genes using multi-dimensional transcriptional comparison, we prioritized identifying novel, sorghum-native developmental regulators associated with key developmental regulatory networks.

To the best of our knowledge, these genotypes have not previously been transformed using either conventional or DR-mediated approaches. Importantly, the molecular pathways that induce SE in monocots remain largely uncharacterized, and genotype-specific responses to morphogenic regulators suggest that the underlying regulatory networks differ among sorghum genotypes.

Through this experimental design, we identified novel transcriptional modules, key regulators, and potential bottlenecks in sorghum transformation, enabling the rational design of improved, more universal transformation systems. Ultimately, this study functionally validated novel genes, provided a framework to enhance transformation efficiency in sorghum, and offered insights that may apply to other monocot crops.

## Materials and methods

### *Agrobacterium tumefaciens* strain and plasmid vectors

For all experiments, *Agrobacterium tumefaciens* Thymidine-auxotrophic strain LBA4404 Thy-containing the helper plasmid ternary vector transformation system pPHP71539 (Anand *et al*., 2018; Zhang *et al*., 2019) was used. For immature-embryo transformation, two vectors were used: (1) the T-DNA plasmid vector pPHP96021 carrying *ZmpltP:ZmWUS* and *CmYLCV:Ruby* as a visible marker. Here, *CmYLCV:Ruby:HspT* was amplified using attB4r and attB3 flanking sites and subsequently cloned into pDonr221 with attP3 and attP4 sites via BP cloning (Roure *et al*., 2007), then cloned into pPHP96021 and pPHP69219 (containing attL4 and attL3 sites) via LR cloning. (2) The GRF:GIF vector containing *ZmUbi:TaGRF4:GIF1:NosT* was amplified from JD633 (Debernardi *et al*., 2020) and cloned into module C. *ZmUbi:Cyan:PinII* (fluorescence marker) was amplified from pPHP81858 and cloned into module B; both modules were combined via one-step Golden Gate assembly into the pTrans220d transformation backbone containing the *neomycin phosphotransferase II* (*NptII*) selectable marker from the Voytas toolkit. The empty control vector consisted of *PvUbi2:GFP* (pModC_3013) and *NptII* (pTrans220d) (Čermák *et al*., 2017). For leaf transformation, pPHP81858 (containing *Nos:WUS2* and *Ubi:BBM* along with *ZmUbi:Cyan* as visible marker) and pPHP69219 (with *Nos:WUS2* and *Ubi:BBM* along with *CmYLCV:Ruby* as visible marker) were used. All vectors included the *NptII* selection marker except pPHP69219 (no selection used for leaf transformation). pPHP96021 employs a heat-shock-inducible Cre-loxP system for excision of *ZmWUS* and the *NptII* marker. Similarly, pPHP81858 and pPHP69219 use a desiccation-inducible Cre-loxP system to excise *Nos:WUS2* and *Ubi:BBM*. In contrast, the GRF-GIF construct was retained without an excision system, as previous studies report that its expression did not cause pleiotropic effects (Butler *et al*., 2025; Li *et al*., 2024). Primer sequences used for construct design are given in File S1; vector maps are provided in **Figure S1.**

### Plant material, explant preparation, and *Agrobacterium* Infection

The study used sorghum inbred line Tx2910 (PI 598070), a Sudangrass-type genotype (Rooney *et al*., 1998), and Tx2783 (PI 656001), a sorghum-aphid-resistant germplasm (Peterson *et al*., 1984). We adapted the immature-embryo (IE) transformation protocol from Che *et al*., 2022. Briefly, we collected immature seeds 13–15 days after pollination from greenhouse-grown plants and stored them overnight at 4°C. We selected uniform-sized seeds, sterilized them in 30% bleach with 0.1% Tween-20, and rinsed them 5–6 times with sterile water. IEs approximately 2-2.5 mm were excised using sterile tweezers and a scalpel; approximately 30–40 IEs were transferred to a 2 mL tube containing infection media and washed 2–3 times to remove milky endosperm. IEs were heat-treated (45°C, 3 min), then incubated in *Agrobacterium* suspension supplemented with 200 µM acetosyringone, 50 mg thymidine, and 10% Pluronic-F68 surfactant (OD550 = 0.4) for 10 min in the dark. Treated embryos were plated onto co-cultivation medium (scutellum side up), sealed with micropore tape, and incubated in the dark at 24°C overnight, followed by 3 days at 28°C (Che *et al*., 2022). For leaf transformation, mature seeds were surface-sterilized with 75% ethanol followed by 30% bleach with Tween-20, then transferred to germination medium supplemented with 2 mg/mL ancymidol. *In vitro* germination was carried out at 28°C with a 16/8 h light/dark photoperiod. Before explant harvesting, seedlings were placed at 45°C and 70% relative humidity for 3 h. Leaf-base segments (∼3 cm) were harvested and fragmented using a Cuisinart Mini-Prep Food Processor. For *Agrobacterium* infection, OD550 = 0.6 was prepared in infection medium supplemented with 200 µM acetosyringone, 50 mg thymidine, and 10% Pluronic-F68 surfactant (Thermo Fisher Scientific, Cat. No. 24040032). Leaf explants were infected for 20 min at room temperature, blotted dry, and transferred to solid co-cultivation medium; plates were incubated at 28°C in the dark for two days (Wang *et al*., 2023b).We based the media composition on published protocols for IE transformation (Che *et al*., 2022) and leaf transformation (Wang *et al*., 2023b) with minor modifications. G418 (250 mg/L) was used to increase selection stringency and minimize escapes.

### Transformation and analysis of transgenic events

To avoid ambiguity from ratios that can exceed 100% when multiple shoots regenerate from a single explant, we used the following efficiency matrix (1) Transformation efficiency (%) = (Number of IEs producing embryogenic structures with visible marker gene expression 7 DPI/ Total number of starting explants) × 100; this value is bounded 0–100% and reflects the proportion of explants that responded. (2) Regeneration index = total number of plantlets recovered ÷ total number of starting explants, reported as a unitless ratio (plantlets per explant) rather than a percentage; because a single embryogenic callus can give rise to more than one shoot, this index is expected to routinely exceed 1 and should not be expressed as a bounded percentage. (3) Transformation frequency or Transgenic purity (%) = (Number of Ruby or AmCyan plantlets with visible expression as well as PCR confirmation for transgene/Total number of regenerated explants) × 100 (**Table S2**). We performed PCR-based genotyping using *Ruby* primers for the pPHP96021 and pPHP69219 constructs, *AmCyan* primers for the pPHP81858 construct, and *GIF*-specific primers for GRF-GIF construct. The list of primers is provided in **File S1**.

### Sample Collection for bulk and single nucleus RNA-seq

For transcriptome analysis, immature embryo samples were collected after *Agrobacterium-*mediated transformation with control or EV (No DR), WUS (pPHP96021+Ruby), and GRF-GIF at 7 DAT (Days After Transformation) and 15 DAT for both genotypes. Leaf samples from the control were collected at 7 DAT as later stages exhibited necrosis in the absence of DRs, while WUS/BBM (pPHP81858) samples were collected at 7 DAT and 20 DAT. These time points correspond to key stages of SE formation and the initiation of regeneration (**Figure S2**). A summary of sample designations is given in **Table S1**. Each sample name corresponds to a specific genotype, explant type, developmental regulator gene used, and timepoint. Samples were flash-frozen upon collection and ground in liquid nitrogen. mRNA was extracted using the Invitrogen PureLink RNA mini kit. We prepared libraries and performed Illumina sequencing following the standard NovaSeq 6000 PE150 protocols. Three biological replicates were collected for each sample. Single-nucleus RNA-seq was performed using sorghum genotype Tx2783 to compare control (Empty vector with *AmCYAN* marker) vs DR treatment (BBM+WUS; pPHP81858) samples. Samples were collected at 7 DAT to compare bulk RNA seq data. Samples were flash-frozen upon collection and ground in liquid nitrogen. Nuclei were isolated using an adapted protocol from D’Agostino et al. 2025. Briefly, frozen tissue was ground in a mortar and pestle in liquid nitrogen. Samples were transferred to nuclei isolation buffer, passed through 100 µm, 70 µm, and 30 µm cell filters (Miltenyi Biotec), and the filtrate was centrifuged to collect the nuclei pellet. The nuclei washed once with wash buffer and spun again. The final washed pellet was resuspended, stained with propidium iodide, and counted with the Luna-FL cell counter (Logos Biosystems). The nuclei were then loaded onto the chip per the manufacturer’s recommendations to target 10,000 recovered nuclei (10X Genomics, Pleasanton, CA, USA). Library construction for Illumina sequencing was performed with the Chromium™ Single Cell 3’ Library & Gel Bead Kit v3.1 protocol (10X Genomics). The single-indexed sequencing and paired-end libraries were carried out on an Illumina™ NovaSeq 6000 platform according to the 10X Genomics manual (D’Agostino *et al*., 2025).

### RNA-seq data analysis

Raw fastq.gz files were processed with fastp (0.23.2) using "–detect_adapter_for_pe," "–trim_poly_x," and "–trim_poly_g" to remove adapters and low-quality reads (Chen *et al*., 2018). We used the *Sorghum bicolor* BTx623 v3.1 genome as reference (McCormick *et al*., 2017). To confirm transgene integration, transgene sequences (*ZmWUS, ZmBBM, Ruby, GRF-GIF, AmCyan*) were added to the reference annotation. Filtered reads were mapped to the modified reference using kallisto (0.48.0) (Bray *et al*., 2016). We used kallisto abundance files for differential expression analysis in R with DESeq2 (1.44.0), applying a threshold of |log2FC| ≥ 2 and Benjamini–Hochberg-adjusted *P* < 0.05 to define DEGs for each pairwise contrast (Love *et al*., 2014). Variance-stabilized transformation (VST) values were used for PCA visualization, and Z-scaled VST values for k-means clustering, visualization, and hormone dot plots. The optimal cluster number for k-means clustering was determined using the elbow and silhouette methods implemented in fviz_nbclust (factoextra 1.0.7), followed by visual inspection of heatmap coherence (Kassambara and Mundt, 2016). Heatmaps were generated using "pheatmap" within ComplexHeatmap (2.20.0) (Gu, 2022). GO enrichment used the topGO package in R with annotations from Phytozome (Goodstein *et al*., 2012; Gu, 2022). Gene regulatory network (GRN) analysis used the "GENIE3" algorithm within the bionero package (1.17.3) (Almeida-Silva and Venancio, 2022); each sample was subset to its top 5,000 expressed genes prior to GRN construction. Gene co-expression network (GCN) analysis used the top ∼10,000 variable genes per explant type (embryo or leaf samples), analyzed via the "WGCNA" algorithm in bionero with *net_type* = "signed hybrid" and a module-merging threshold of 0.7. GRNs and GCNs were visualized in Cytoscape (3.10.2) (Shannon *et al*., 2003). Hub TFs were defined separately for the two network types used in this study, since they capture conceptually different properties. For GRN analysis, hub TFs were defined as the top 10% of regulatory nodes within each expression module by weighted out-degree (predicted regulatory connections to target genes). For GCN analysis, hub genes were defined using standard WGCNA criteria: top 10% of genes based on module membership (kME) as well as those with a kME > 0.8.

### snRNA-seq data analysis

Sequencing data was processed using kallisto bustools to obtain cell feature counts for analysis (Sullivan *et al*., 2025). To confirm transgene integration, transgene sequences (*ZmWUS, ZmBBM, AmCyan*) were added to the reference annotation. Count data was further analyzed within the R package Seurat v5. (Hao *et al*., 2024). Nuclei in the 95th percentile for unique gene count and 90^th^-99^th^ percentile for UMI were selected for further pre-processing. Putative doublets were removed with scDblFinder, and background RNA was removed with SoupX (Germain *et al*., 2022). The data were then normalized by SCTransform and processed via runPCA. Samples were aligned with align_cds and reducedDim, both from the Monocle 3 R package, and UMAP plots were created; clusters were assigned via cluster_cells (Cao *et al*., 2019). Annotation of cell types/states was performed by utilizing orthologs from published snRNA-seq datasets from soybean seed, SAM and rice regenerative callus (Lyu *et al*., 2026) (**File S15**), highlighting select marker genes within clusters (Seurat FindAllMarkers with min.pct = 0.3, logfc.threshold = 0.3, min.diff.pct = 0.2) and GO term enrichment analysis. Expression of marker gene sets were analyzed using the AddModuleScore function from Seurat and observed using the dotplot and featureplot functions. GO term enrichment was performed using the topGO package in R (Alexa and Rahnenfuhrer, 2010) with gene annotations from the *Sorghum bicolor* BTx623 v3.1 genome obtained from Phytozome. Trajectory and pseudo-time analysis were performed using Monocle3, and other plots were visualized using ggplot2 (Gómez-Rubio, 2017).

### Validation of the novel DR genes

The coding sequences of the candidate genes were either PCR-amplified from the cDNA or commercially synthesized (Twist Bioscience), including the attB1 and attB2 sites. We codon-optimized several genes to facilitate synthesis while preserving the native amino acid sequence. The coding sequences were first cloned into the pDONR221 Zeocin entry vector using the Gateway® BP recombination reaction. Subsequently, the genes were transferred into the Gateway-compatible binary vector pANIC6A by an LR recombination reaction placing each gene under the control of the maize ubiquitin (*ZmUbi*) promoter for constitutive expression (Mann *et al*., 2012). We performed *Agrobacterium*-mediated transformation using the Thy-LBA4404 strain, along with a helper plasmid, and regenerated plants in Tx2783 genotypes, as described above. For preliminary experiments, we used the pANIC6A overexpression vector with candidate genes cloned for transformation. Candidate genes that showed a positive embryogenic response in preliminary experiments were subsequently transferred to the pMOD_A vector (*ZmUBI:GENE:OcsT*) and cloned with the *CmYLCV:AmCYAN* fluorescent marker (pMOD_C) using golden gate assembly (Čermák *et al*., 2017; Devkar *et al*., 2025) to facilitate rapid identification of transformed tissues. *ZmWUS* + *Ruby* (pPHP96021) or *ZmWUS* + *AmCYAN* (pPHP96564), and ZmWUS+ZmBBM (pPHP81858) were included as positive controls.

### Morphological imaging and histological analysis of somatic embryos

The progression of SE formation was monitored at different time points (7, 12, 15, 20 DAT) by morphological imaging using Olympus SZX10 stereo microscope. Images were captured under comparable magnification and illumination conditions. For scanning electron microscopy (SEM), samples were freeze-dried and mounted on double-sided carbon tape. We sputter-coated samples with iridium using a Safematic CCU-010 compact coating unit and imaged them with a Hitachi S3400 SEM (Hitachi, Ltd., Tokyo, Japan) (Dhiman *et al*., 2025). Images were acquired at a working distance of 15 mm and 15 kV beam acceleration at different magnifications. For histological analysis, developing calli with SE were fixed in FAA solution containing 90 mL of 50% ethanol, 5 mL of 37–40% formaldehyde, and 5 mL of glacial acetic acid. Samples were fixed for 12–24 h at room temperature, followed by dehydration through a graded ethanol series and clearing with xylene. Samples were then infiltrated with paraffin and embedded for sectioning. Paraffin-embedded samples were sectioned at 5–8 µm thickness and mounted on glass slides. Sections were deparaffinized with xylene, rehydrated through a descending ethanol series to water, and stained with Toluidine Blue O. Following staining, sections were rinsed, dehydrated through an ethanol series, cleared with xylene, and used for microscopic examination (Steinmacher *et al*., 2007).

## RESULTS

### DRs enabled efficient transformation and regeneration in sorghum genotypes

We selected the grain sorghum inbred line Tx2783 and the forage sorghum line Tx2910, neither of which had previously been evaluated for tissue-culture response or regeneration capacity. IEs of both genotypes were transformed with either an empty vector or a construct expressing DR genes (WUS2 or GRF-GIF). In parallel, leaf segments were transformed with a WUS2 and BBM combination. Transformation with the empty vector rarely to produce embryogenic calli and failed to regenerate in either genotype across two independent experiments, confirming their recalcitrant nature under conventional methods. In contrast, DR-mediated transformation markedly enhanced regeneration efficiency in both genotypes (**Figures 1–3**). We first evaluated the *WUS2* construct pPHP96021 using *Ruby* as a visual marker (**Figure 1**). In Tx2910, 70% of explants produced embryogenic calli and we regenerated 108 plantlets in total, 86 of which showed red shoots due to Ruby marker expression (regeneration index=2.7 plantlets/explant), confirming successful transgene incorperation. In Tx2783, 37% of explants produced embryogenic calli, generating 188 plantlets in total, of which 169 were Ruby-positive (Regeneration index=0.94) (**Figure 1 A–C, Table S2**). No DR-mediated pleiotropic effects were observed in the regenerating calli due to heat-shock-promoter-driven Cre-loxP-mediated excision of the DR (*WUS2*). Although *Ruby* expression was highly observable in callus and regenerated plantlets (**Figure 1B, C**), it’s presence gradually diminished in the vegetative tissues as the plants developed, but was consistently observed in the leaf base, reproductive tissues, and seeds (**Figure S3A–F**). This expression pattern is consistent with earlier reports that the *CmYLCV* promoter is highly active in meristematic and reproductive tissues (Kshetry *et al*., 2025; Stavolone *et al*., 2003). Overall, these results indicate that *WUS2* can efficiently transform and regenerate both genotypes.

**Figure 1.**
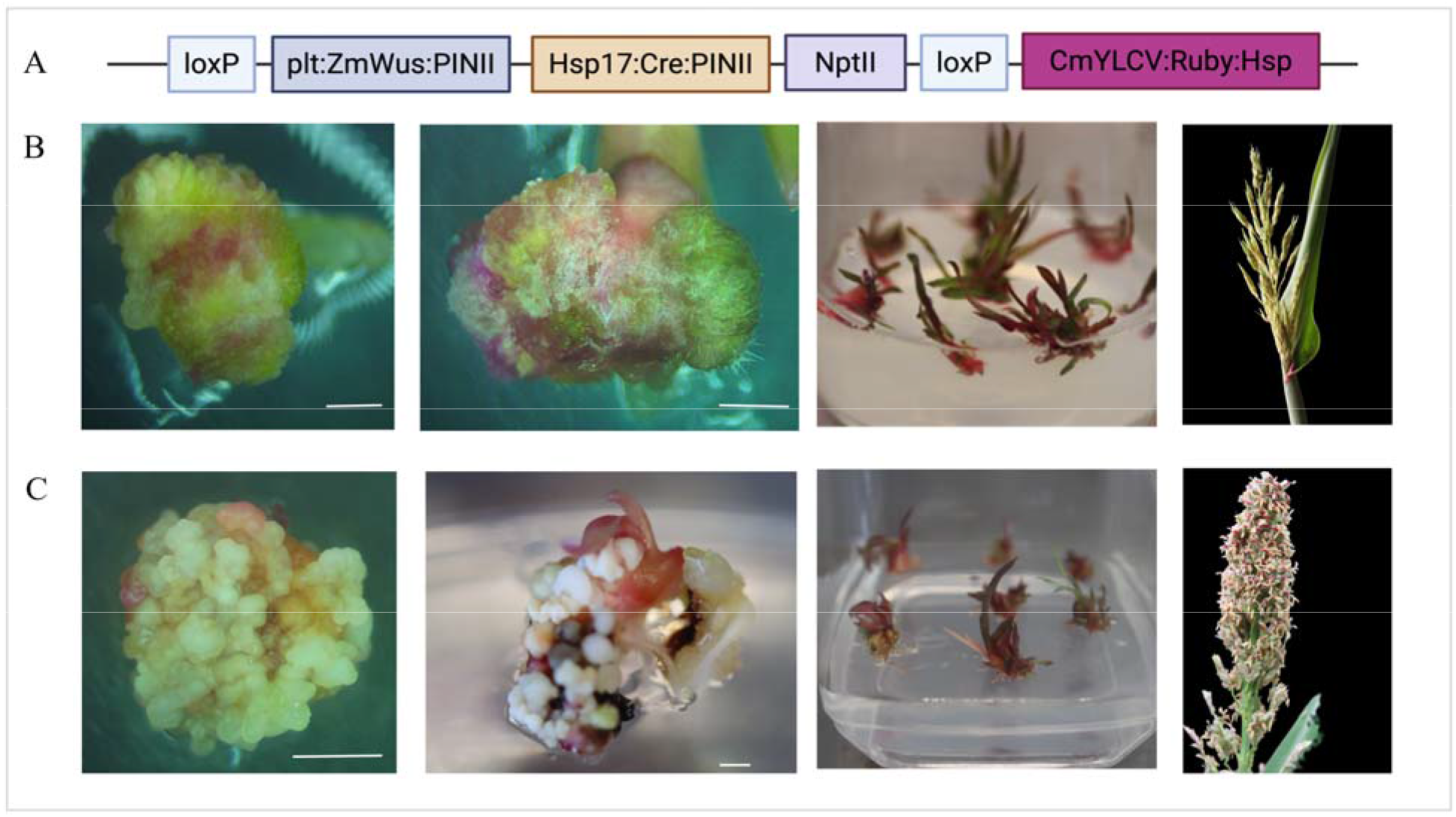
Schematic representation and phenotypic outcomes of sorghum immature embryo transformation using ***ZmWUS***.(A) T-DNA vector design for *Agrobacterium*-mediated transformation. (B) *ZmWUS*-mediated transformation in forage sorghum (Tx2910). (C) *ZmWUS*-mediated transformation in grain sorghum (Tx2783), showing *Ruby* expression in callus, regenerated plantlets, and panicles. Scale bar = 3 mm.

**Figure 2.**
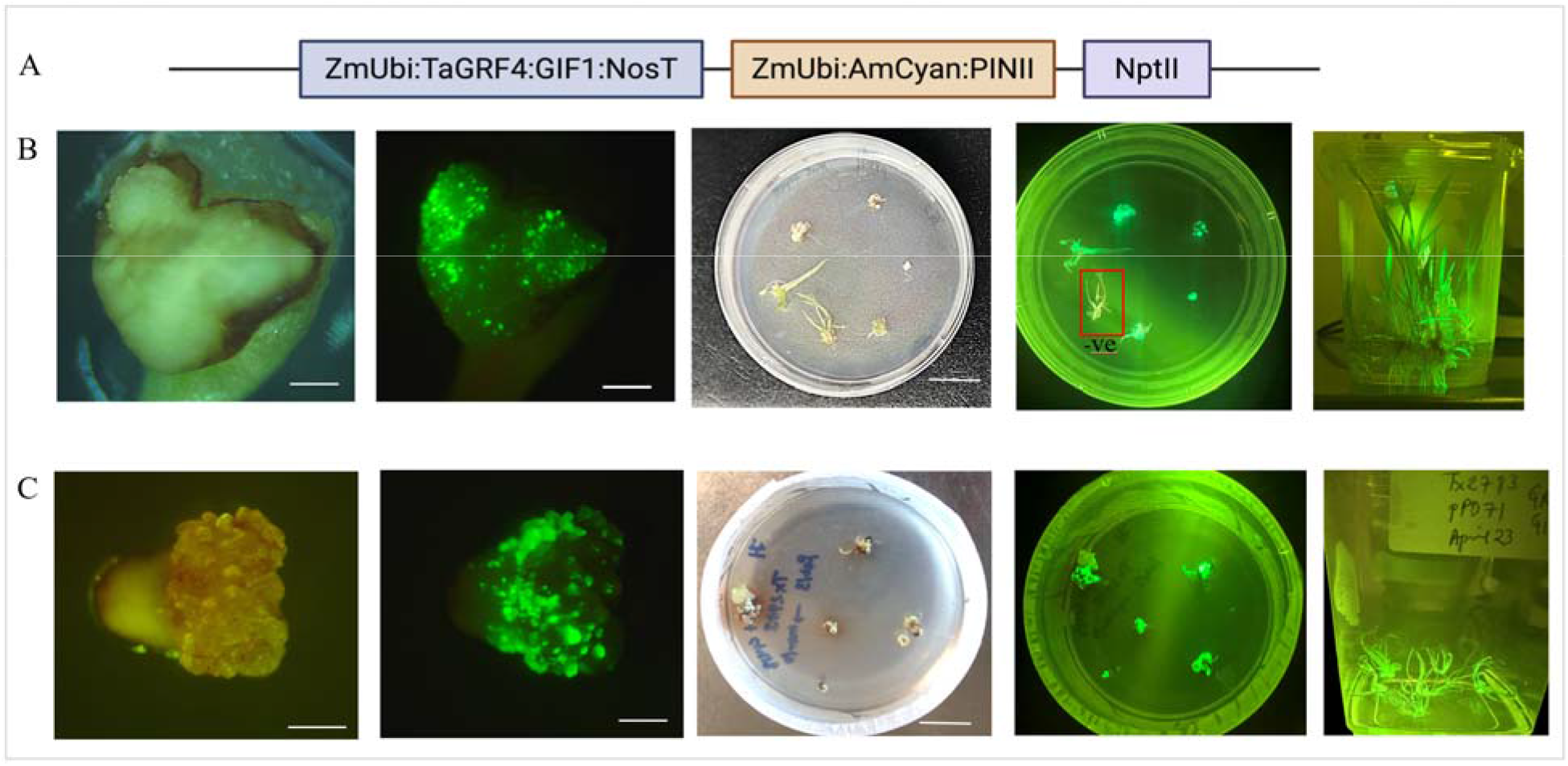
Schematic representation and phenotypic outcomes of sorghum immature embryo transformation using ***TaGRF-GIF*** chimera. (A) T-DNA vector design for *Agrobacterium*-mediated transformation. (B) GRF-GIF-mediated transformation in forage sorghum (Tx2910). (C) GRF-GIF-mediated transformation in grain sorghum (Tx2783). Fluorescent somatic embryos expressing *AmCyan* were observed 5–7 DAT, followed by plant regeneration. Scale bar = 3 mm.

**Figure 3.**
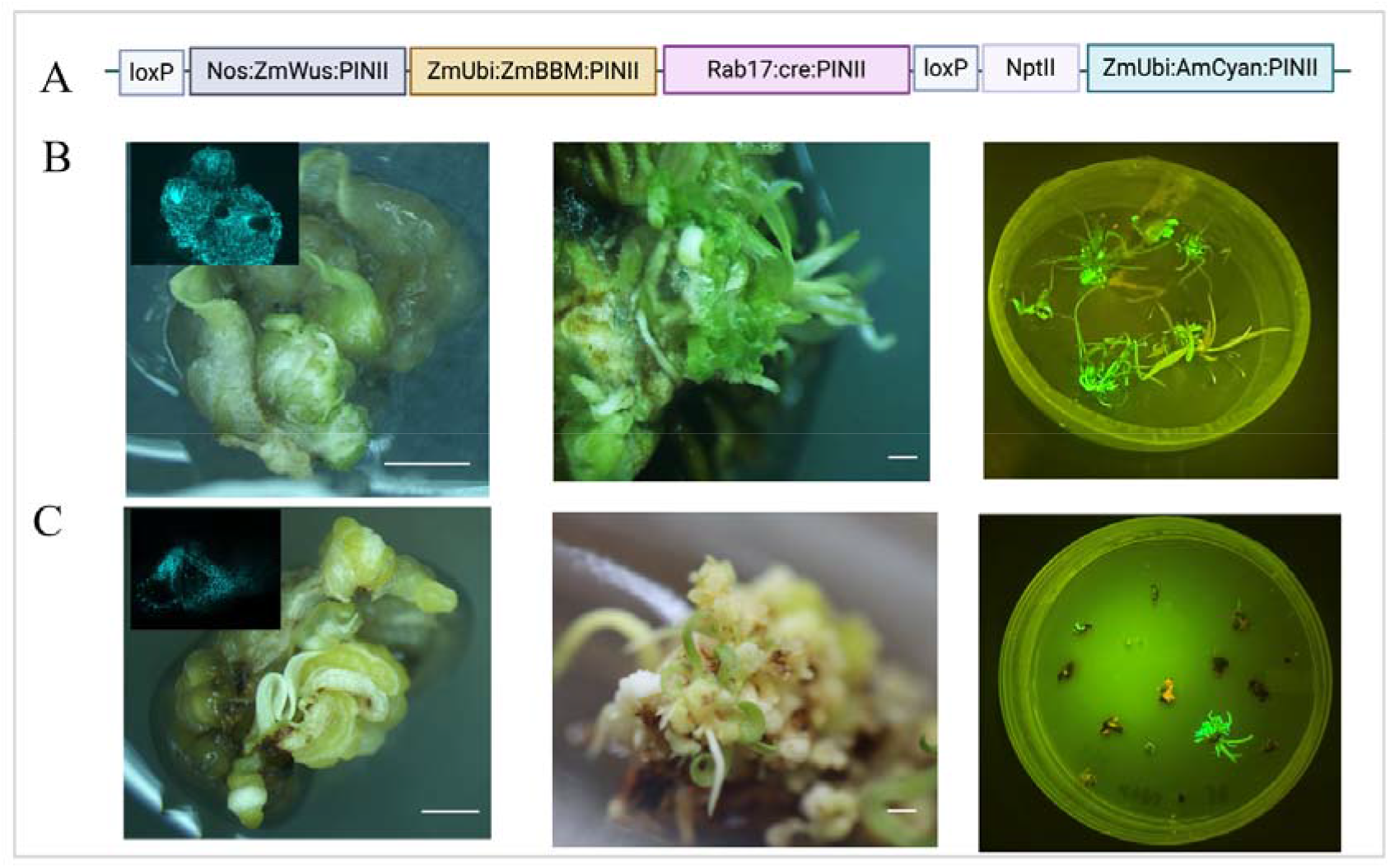
Schematic representation and phenotypic outcomes of sorghum leaf base transformation. **(A)** T-DNA vector design used for transformation. **(B)** Somatic embryogenesis and plant regeneration in forage (Tx2910) from leaf base explant (**C**) Somatic embryogenesis and plant regeneration in grain sorghum (Tx2783) from leaf base explant, showing *AmCyan* fluorescence in callus and regenerated plantlets. Scale bar = 3 mm.

To further assess an alternative DR, we tested the *TaGRF-GIF* chimera driven by the *ZmUbi* promoter, which has been shown to drive efficient transformation in wheat (Debernardi *et al*., 2020), along with *AmCyan* as a visual reporter (**Figure 2A–C**). In Tx2910, 38 of 75 IEs (51%) formed embryogenic calli, yielding 110 plantlets (regeneration index = 1.47 plantlets/explant), 84 (76%) of which were AmCyan-positive. In Tx2783, 32 of 65 IEs (49%) formed embryogenic calli, giving rise to 84 plantlets (regeneration index = 1.29 plantlets/explant), of which 68 were AmCyan-positive (**Table S2**). Overall, the lesser embryogenic calli response in Tx2783 with *WUS2* and *GRF-GIF* compared to Tx2910 was associated with pronounced phenolic exudation (**Figure S4**). At the seed-setting stage, all T0 plants with constitutive *GRF-GIF* chimera expression under the *ZmUbi* promoter showed a substantial reduction in seed set. The effect was particularly severe in Tx2910, where seed number was reduced to as few as 0–5 seeds per panicle (**Figure S5A, B**).

We further explored leaf segments as an explant source by transforming Tx2910 and Tx2783 *with Nos:WUS2* and *Ubi:BBM* co-expressing *AmCyan* (**Figure 3A**). This promoter and DR combination had been previously used in by Lowe et al. 2016 for maize transformation, which produced embryogenic callus response (Lowe *et al*., 2016). The same DR combination, but with enhanced promoters, was previously used for sorghum leaf-base transformation in Tx430, giving rapid somatic-embryo formation (Wang *et al*., 2023b) and more recently for gene editing and transformation across multiple sorghum genotypes (Azanu *et al*., 2025). Although SE from leaf explants using *Nos:WUS2* and *Ubi:BBM* was comparatively slower than from IEs, we observed embryogenesis within 7–10 days after transformation and recovered fully regenerated plants within approximately 3–3.5 months (**Figure 3**). *AmCyan* fluorescence and further molecular analysis confirmed transgene integration in regenerated plantlets (**Figure 3B, C; Figure S6, S7C, D**).

In a separate experiment using leaf-base explants with the same DR combination and *Ruby* as a visual marker under a selection-free strategy, we obtained reproducible transformation results (**Figure S8**). Because constitutive *WUS and BBM* expression causes pleiotropic effects, these cassettes were flanked with a CRE-loxP excision system. As previously observed, whenever *WUS/BBM* failed to excise, pleiotropic effects appeared in Tx2783 (**Figure S5C, D**). We also tested the *GRF-GIF* chimera for leaf-based transformation, but it failed to induce regeneration in either genotype. **Table S2** summarizes the transformation outcomes, including the number of initial explants, transformation, and regeneration efficiency. **Figure S3G, H,** and **S6A-D** show T1 segregating individuals from both genotypes with *Ruby* and *AmCyan* phenotypes, respectively. Collectively, these results show that forage type Tx2910 has superior tissue culture responsiveness with minimal phenolic interference, despite a higher escape frequency under G418 selection. The elevated escape rate may reflect forage sorghum’s intrinsic regenerative vigor and tillering capacity and/or transient or non-cell-autonomous expression of DR genes in certain cells, leading to shoot regeneration without transgene integration.

### DRs reprogram transcriptional networks and promote regeneration competence

Although transformation outcomes varied among genotypes, overall efficient regeneration by DR overexpression suggested a role in reshaping developmental responses. To investigate the underlying molecular basis, we compared transcriptomic profiles of DR-expressing and control tissues during early embryogenesis and regeneration competence acquisition in the two sorghum genotypes (**Table S1**). We selected embryogenic calli at 7 and 15 DAT from immature-embryo explants across control, *GRF-GIF*, and *WUS* overexpression treatments. For leaf explants, we selected control samples at 7 DAT and *BBM+WUS2* overexpression samples at 7 and 20 DAT. The replicates showed a high correlation with an average R^2^>=0.9. Following the replicates, QC samples with detectable transcripts from at least one of the transgenes from the vector used (*ZmBBM*, *ZmWUS2*, *GRF-GIF*, *Ruby*, *AmCyan*) were retained for downstream analysis. (**File S2**). Differential expression analysis revealed genotype-specific responses as well as conserved DR-responsive programs (**Table S3; Figures S9, S10**). Both *WUS* and *GRF–GIF* induced progressively larger transcriptional changes from 7 to 15 DAT, with overlapping genotype-independent responses enriched for pathways associated with cellular development, metabolism, photosynthesis, hormone regulation, sucrose metabolism, and cell-wall remodeling.

This conserved reprogramming is reflected in 387 and 604 overlapping differentially expressed genes (DEGs) between genotypes at 7 and 15 DAT for *WUS* (**Figures S9**), and from 20 shared DEGs at 7 DAT to 339 at 15 DAT for *GRF-GIF* (**Figures S10**). The shift from a comparatively modest early response (7 DAT) to a larger response at 15 DAT indicates that these DRs act primarily by reinforcing transcriptional programs required to establish regeneration competence. Transcriptome analysis of leaf-derived embryogenic callus further demonstrated strong genotype-and time-dependent responses following *WUS+BBM* expression. At 7 DAT, FL_BW (F= forage type, Tx2910 background) showed 8,151 upregulated genes relative to control. Temporal comparison (7 vs. 20 DAT) showed 8,481 genes upregulated at 7 DAT versus 6,535 at 20 DAT. GL_BW (G=grain type, Tx2783 background) showed 4,895 upregulated genes at D7 relative to control, and 4,991 vs. 3,330 upregulated genes in the 7 vs. 20 DAT comparison (**Table S3**). The strong WUS+BBM-induced upregulation in leaf-base callus reinforces the role of combinatorial DR activity in driving embryogenic reprogramming, with genotype-dependent magnitude differences.

### *GRF-GIF* and *WUS* accelerate SE in immature embryos through distinct transcriptional modulations

K-means clustering of DEGs coupled with GO (Gene Ontology) enrichment revealed coordinated, regulator-and genotype-specific transcriptional shifts underlying SE (**Figures 4, 5)**. In forage sorghum Tx2910, six co-expression clusters captured this reprogramming (**Figure 4A**). Cluster 1, associated with *WUS* treatment was enriched in GO pathways promoting cell-fate identity, and mobilizing metabolic reserves for dedifferentiation. Cluster 5 (all 7 DAT samples) showed enrichment for cell-fate specification, dormancy, and catabolic regulation, consistent with an early, transient suppression of differentiation. Clusters 2 and 3 (control_D15, GIF_D15) were enriched for culture induced stress and growth-related GO terms; their downregulation specifically in WUS-expressing tissue indicates that WUS actively suppresses these persistent stress/cell-wall-remodeling programs. Cluster 6, induced in D15 samples, represents a developmental-competence module (cell communication, signaling, meristem/reproductive-identity specification) that sustains fate-specification programs needed for regeneration (**Figure 4B; Files S3, S4**). Similarly, in grain sorghum, among four K-means clusters, Cluster 1 (abiotic stress, secondary-metabolite/phenolic pathways) was high in control and GRF-GIF_D7 but strongly downregulated in WUS_D7, WUS_D15, and GIF_D15, indicating that WUS helps reprogram stress-dominant states as early as 7 DAT, while GRF-GIF’s suppressive effect is delayed until 15 DAT (**Figure 5A, B**). Cluster 2 (cell/organ morphogenesis) was induced in WUS (D7, D15) and GRF-GIF_D15, and Cluster 4 contained genes expressed in all conditions except control_D15 indicating that, without DRs, Tx2783 remains locked in a stress-dominant state driven by high phenolic release. Overall, grain type Tx2783 showed a greater dependence on DR-mediated activation of developmental pathways than forage type Tx2910.

**Figure 4.**
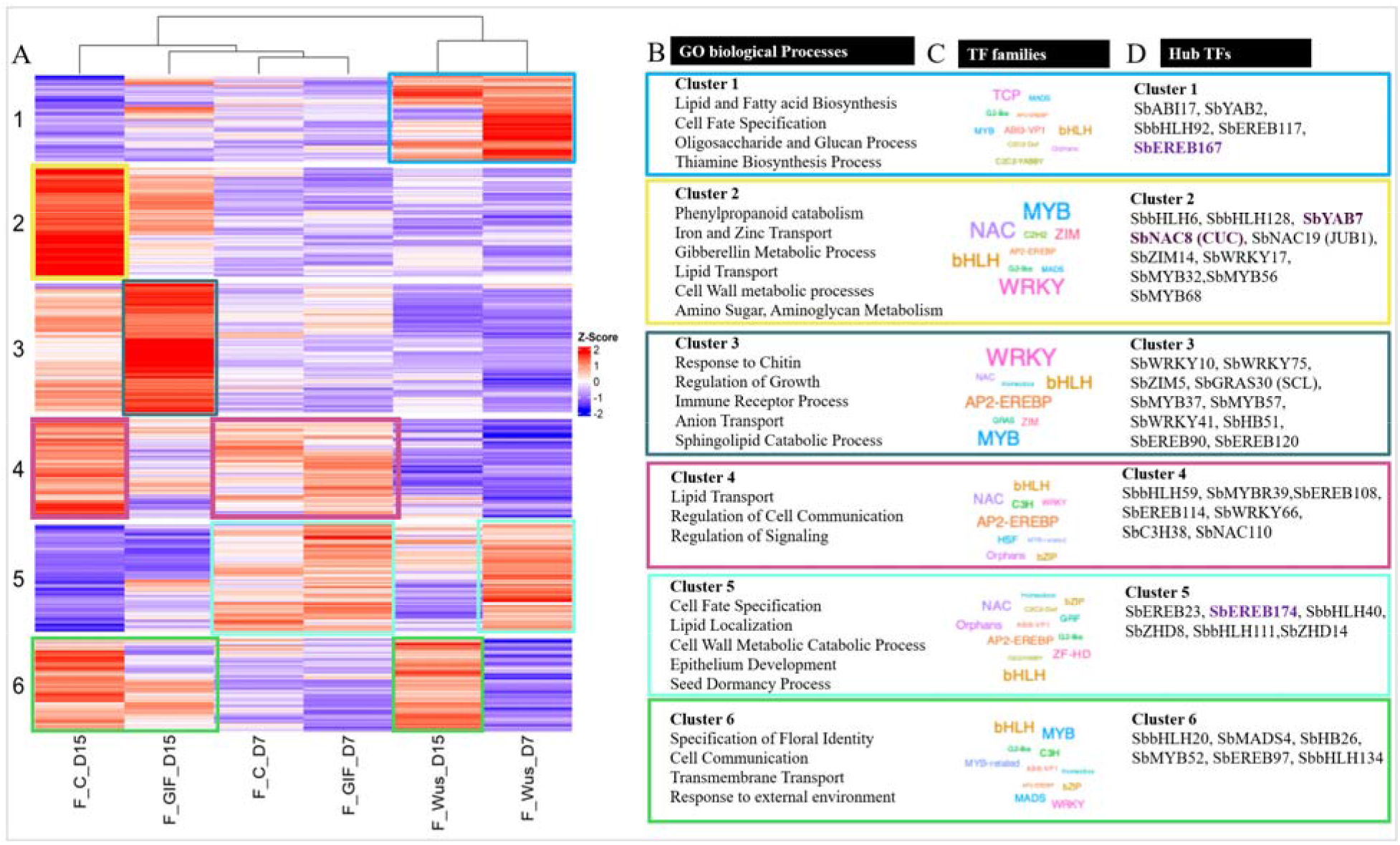
K-means clustering-based identification of transcriptional modules responding to a developmental regulator in forage sorghum (Tx2910). (A) K-means clustering heatmaps of differentially expressed genes (log2FC >=2, an adjusted p-value (Benjamini–Hochberg corrected <0.05) across samples (control, GRF-GIF, and WUS) at 7 and 15 days after transformation (DAT) in Tx2910 (B) Representative enriched Gene Ontology (GO) biological process terms (C) TF family frequency within each cluster; Text size is proportional to the number of TF family members within each cluster, larger text indicates higher counts. (D) Representative hub transcription factors (TFs) from each cluster/sample (with known roles in development, hormone related, pathogen interaction, flavonoids, cell proliferation, etc.).

**Figure 5.**
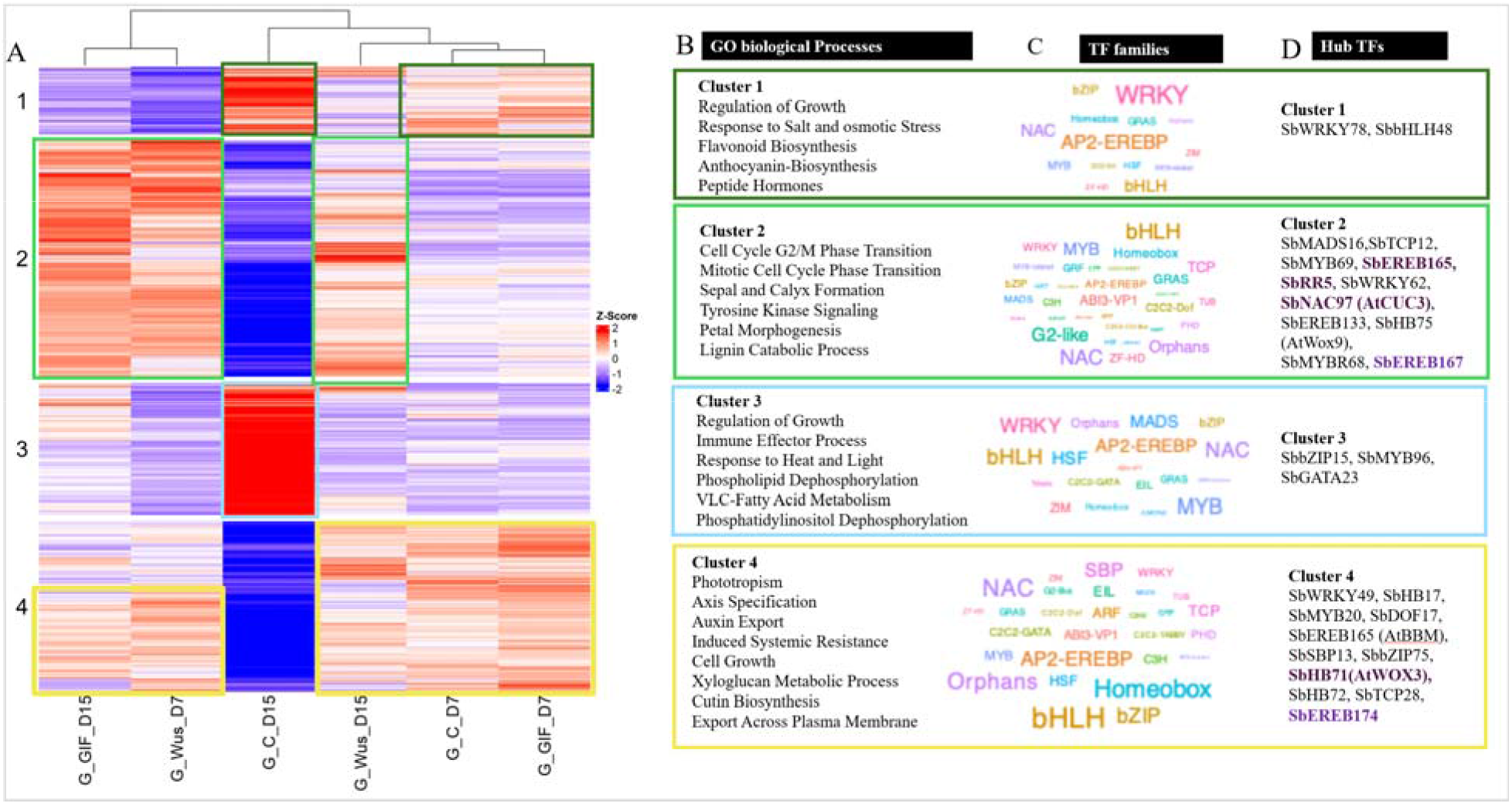
K-means clustering-based identification of transcriptional modules responding to a developmental regulator in grain sorghum (Tx2783). (A) K-means clustering heatmaps of differentially expressed genes (log2FC >=2, an adjusted p-value (Benjamini–Hochberg corrected<0.05) across control, GIF, and WUS treatments at 7 and 15 days after transformation (DAT) are shown in Tx2783 (B) Representative enriched Gene Ontology (GO) biological process terms (C) TF family frequency within each cluster; Text size is proportional to the number of TF family members within each cluster; larger text indicates higher counts. (D) Representative hub transcription factors (TFs) from each cluster/sample (with known roles in development, hormone related, pathogen interaction, flavonoids, cell proliferation, etc.).

Further, gene regulatory network analysis of cluster-specific hub TFs revealed both shared and genotype-distinct regulatory architectures underlying developmental control, embryogenic competence, and regeneration potential. In Tx2910, hub TFs from *SbWRKY*, *SbMYB*, *SbNAC*, *SbEREBP/AP2*, and *SbbHLH* families were present in multiple clusters, reflecting probable roles in early morphogenetic reprogramming, hormone signaling, and embryogenesis (**Figure 4 C, D**). For example, a homolog of Arabidopsis CUP-SHAPED COTYLEDON3 (CUC3), *SbNAC8* a hub in F_GIF_D15 is a NAC-domain transcription factor involved in developmental patterning, organ boundary specification, and shoot meristem formation (Hibara *et al*., 2006; Takada *et al*., 2001; Takeda *et al*., 2011). Similarly, *SbbHLH111*, a hub in GRF-GIF samples and homolog of *AtbHLH137*, functions in cytokinin signaling to promote cell proliferation and expansion (Park *et al*., 2021) (**Figure 4D, Files S5, S6**).

Similarly, the grain sorghum Tx2783 regulatory clusters (Clusters 1–4) were dominated by *SbMYB, SbWRKY, SbHB, SbDOF, SbbZIP, SbGATA*, and *SbNAC* families, reflecting genotype-dependent transcriptional control of regeneration (**Figure 5 C, D; Files S5, S7**). Notably *SbEREB165*, the closest sorghum homolog of *ZmBBM* (*Zm00001d042492*), was identified as a hub gene in G_GIF_D15, G_GIF_D7, and G_WUS_D15. *SbNAC97*, a hub in G_GIF_D15 (Cluster 2), is an ortholog of *AtCUC3* and contributes to shoot apical meristem formation (Hibara *et al*., 2006). Similarly, *SbHB75* is a hub in G_GIF_D15, and *SbHB71*, a hub in Cluster 4, both belong to the WUSCHEL-RELATED HOMEOBOX (WOX) family and are associated with meristematic development (**Files S5, S7**). The recurrence of WRKY, MYB, NAC, and EREB hubs across both genotypes underscores their broadly conserved role in embryogenesis and regeneration, while genotype-specific enrichment of DOF, HB, bZIP, and GATA families in Tx2783 points to additional layers of regulatory control governing its more recalcitrant regeneration response. We next compared the ten most highly expressed TFs from four major TF families (AP2-EREBP, ARF, C2C2-Dof, and GRF) comprising 40 TFs in total. Notably, 36 of the 40 top-ranked TFs were shared between genotypes, indicating a largely conserved regulatory core with varying expression patterns. For instance, *SbDOF15, SbEREB174, SbARF26*, and *SbARF13* were uniquely enriched among top TFs in Tx2910, whereas *SbEREB29, SbARF11, SbDOF8*, and *SbDOF12* were specific to Tx2783 (**Figure S11A, B**). These findings enriched our understanding of genotype-specific transcriptional mechanisms and identify candidate TFs for targeted manipulation to improve regeneration efficiency.

### Transcriptional networks driving BBM/WUS2-induced SE

K-means clustering of leaf-derived DEGs (control vs. *BBM+WUS* at 7 and 20 DAT) identified five transcriptional modules in Tx2910 (**Figure S12**) and six in Tx2783 (**Figure S13**). Overall, K-means clustering and GO enrichment in both genotypes indicate that BBM/WUS drives a coordinated transition from stress-and differentiated-tissue-associated programs toward developmental reprogramming; early responses (7 DAT) were dominated by stress signaling and extensive hormone-mediated reprogramming, while later stages were dominated by cell proliferation, morphogenesis, and metabolic activation, reflecting establishment of regenerative competence (**Figures S12B, S13B**). Cluster 3 shows a highly conserved, strongly upregulated profile across both genotypes, with core biological functions related to cell proliferation and structural tissue differentiation, including shared hub genes *SbGRF1* and *SbGRAS74* (**Figures S12C, S13C, File S8, S9**). We next performed weighted gene co-expression network analysis (WGCNA), identifying 18 shared and genotype-specific co-expression modules (41–2,760 genes; **File S10; Figure S14**). Module-trait correlations identified three categories of co-expression programs: (1) BBM/WUS-activated modules, (2) sample-specific early-response modules, and (3) control-state modules antagonistic to DR-driven reprogramming (**File S11**). For instance, Tx2783 (GL_BW_D20) showed a strong association with module 17 (MEblue4), enriched for auxin transport, meristem maintenance, and cell-cycle regulators, consistent with sustained reprogramming activity. Similarly, the Tx2910 (FL_BW_D20) MEblue module showed GO enrichment for functional categories associated primarily with root development and morphogenesis, post-embryonic root morphogenesis (**Figure S14, File S12**). Tx2783 exhibited a stronger transcriptional transition, with activation of DNA replication, RNA processing, ribosome biogenesis, and chromosome organization by 20 DAT (**Figure S14F)**, indicating that the recalcitrant genotype requires more extensive molecular reprogramming to achieve regeneration as evident by K-mean clustering GO analysis and GO ontology of WGCNA module for D20 samples. Furthermore, among AP2-EREB TFs, *SbEREB174* showed the highest expression at 20 DAT in both genotypes. Similarly, *SbARF5, SbARF3, SbARF13*, and *SbARF22* showed elevated expression at 20 DAT DR samples, indicating activation of auxin-responsive transcriptional programs during acquisition of regenerative potential. *SbGRF1* and *SbGRF3* showed marked increases in FL_BW_D20 samples relative to control tissue (**File S13**). Across both genotypes, suppressing defense-and promoting differentiation-associated modules is a conserved prerequisite for regeneration.

### Single-nucleus transcriptomic profiling reveals regeneration-associated cell states

To test whether bulk RNA-seq-identified developmental trajectories and putative regeneration genes can be resolved at the single-cell resolution, we performed single-nucleus RNA-seq (snRNA-seq) on relatively recalcitrant Tx2783 callus at 7 DAT, comparing the empty vector (expressing *AmCYAN* only) and BBM+WUS overexpression (**Figure 6A**). We obtained 3,761 and 3,298 high-quality nuclei from control and transgenic callus, respectively (median 839 UMIs and 606 genes per nucleus), detecting 26,599 of the 34,120 annotated protein-coding genes in sorghum (**Table S4**). Unsupervised clustering identified 21 distinct clusters with strongly skewed composition between samples; cluster 16 for example comprised entirely of cells from transgenic calli (**Figure 6B, Figure S15A, B**). Pseudo time trajectory inference placed these clusters along a continuum from a quiescent, defense-associated state to two proliferative, transgene-associated endpoints. The trajectory originated in clusters 4, 7, and 15 ("Wound-Responsive Embryo-Axis"), marked by elevated general defense and cell-wall-remodeling transcripts and G1/G0 cell-cycle markers (**Figure 6 B, C, S15 D, E**). This cell population reflects the counterpart of the defense-dominant transcriptional state of our bulk k-means analysis identified as the default, DR-suppressed baseline in Tx2783 (**Figure 5 K-mean Cluster 1**). This same signature appeared as a down-weighted expression module in bulk tissue, and here, it appears as a discrete cell population that DR-treated tissue actively moves away from, rather than a uniform tissue-wide change. Clusters 1 and 5 ("Early Embryo-Axis Callus") expressed markers consistent with a transition out of a reserve-mobilizing state, including the rice scutellar-epithelium marker *alphaAmy3/RAmy3D* (Toyofuku *et al*., 1998) (**Figure S17**).

**Figure 6.**
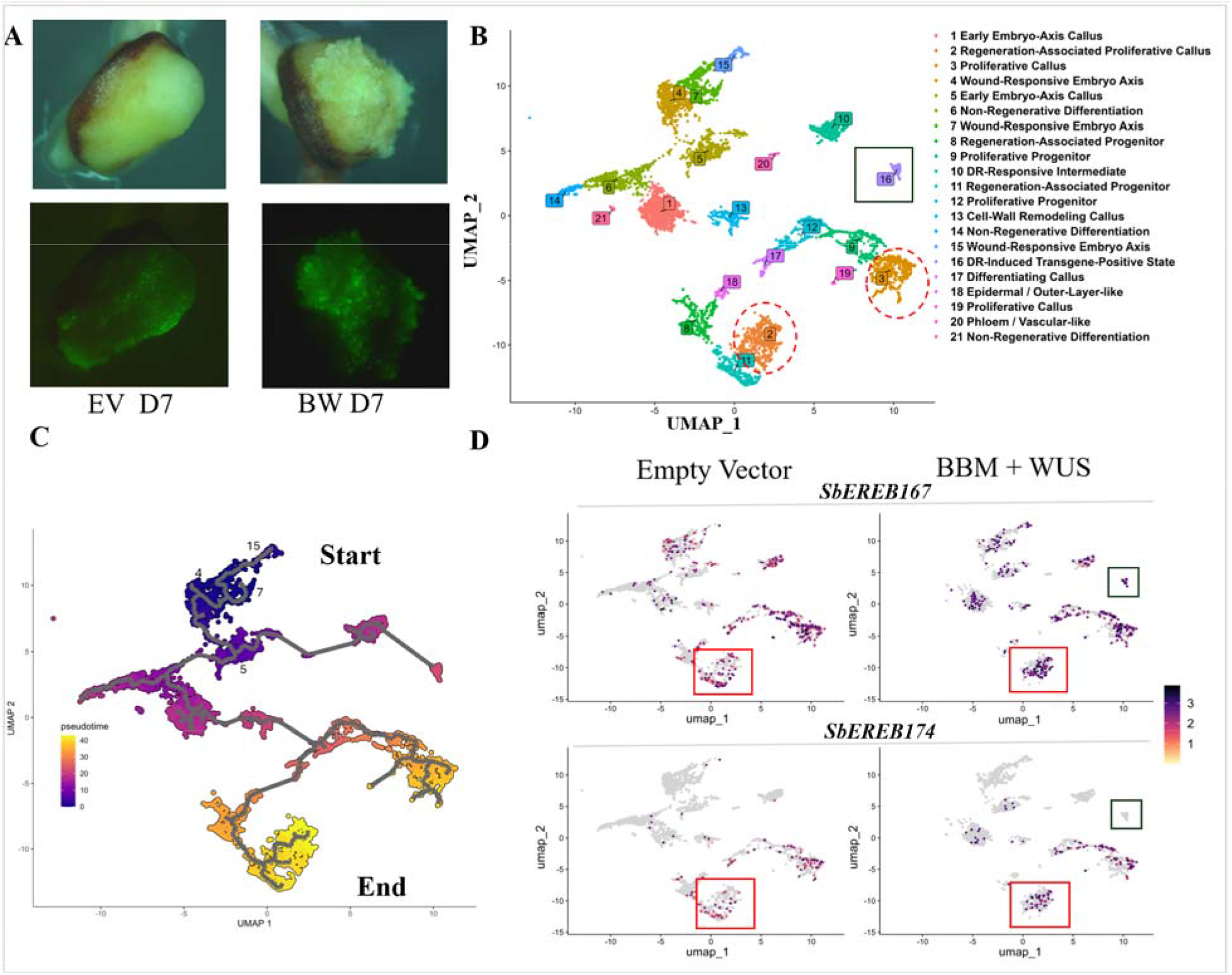
snRNA-seq analysis. **(A)** Representative images of samples collected for nuclei isolation (EV-Empty vector with AmCYAN), BW (BBM + WUS over expression with AmCYAN). **(B)** Integrated UMAP projection of 7,059 high-quality nuclei annotated to 21 differentiated cell states. The color of the cluster represents a cell state as represented in the legend with a dotted circle highlighting the putative proliferative calli. Black open box represents cluster 16 specific to BBM + WUS. **(C)** UMAP plot depicting projected pseudo time of the samples (color represents pseudo time and line indicates pseudo time trajectory). **(D)** UMAP plots depicting localized expression of candidate genes *SbEREB167* (*Sobic.006G245500*) and *SbEREB174* (*Sobic.010G215100)*. The top two plots represent *SbEREB167,* and the bottom represents *SbEREB174*. Red box represents expression in putative proliferative calli. Cluster 16 (black box) specific to BBM + WUS. EV samples are present on the left of each row and BW on the right.

SE from immature-embryo explants in cereals is well studied and it originates from the scutellum rather than from the embryo axis proper (Eudes *et al*., 2006; Vasil *et al*., 1985). The presence of scutellar markers in the earliest founder-like population was notable and suggests the possibility that this cluster is at least partly scutellum-derived. From this founder-like state, the trajectory bifurcated into two proliferative branches marked by S-phase and G2/M cell-cycle activation, wherein cluster 2 and 3 ("Regeneration-Associated Proliferative Callus") were identified using converging markers for "initial/proliferating cells" and "shoot meristem niche" from rice callus snRNA-seq and "SAM" identity from soybean (**Figure S15-S17, File S14, S15**). These proliferative endpoints correspond closely to the cell-cycle module our bulk analysis identified as the hallmark of DR-driven reprogramming in Tx2783 (**Cluster 2, Figure 5: "cell-cycle G2/M phase transition"**), in which *SbEREB167* and *SbEREB165* were identified as network hubs.

Clusters 6 and 14 lacked proliferative/cell-cycle markers and were dominated by non-transgenic nuclei (**Figure S15 A-C**). Cluster 14 additionally showed mesophyll/photosynthetic markers, consistent with untransformed cells and defaulting to a differentiated "greening" callus rather than regeneration (**Figure S16, S17**). Remaining clusters were annotated as "Cell-Wall Remodeling Callus" (cluster 13; *SbEXPB2* orthologs, *SbLRX4*, *SbCWINV2*), "Epidermal/Outer-Layer-like" (cluster 18), and "Phloem/Vascular-like" (cluster 20) (**File S14, Figure S16**).

*SbEREB167* and *SbEREB174* (selected for functional validation below) were detectable broadly across cell types and treatments, including in control tissue, but showed markedly elevated expression specifically within the proliferative/regeneration-associated population under BBM+WUS treatment (**Figure 6D**). This pattern of broad baseline expression suggested that these genes’ activity is gated less by where they are expressed than by how highly they are expressed within a cell already primed for proliferation.

### Novel developmental regulators to promote sorghum embryogenesis

Integrative analysis of DEGs and gene regulatory networks converged on a set of high-confidence candidate genes potentially orchestrating SE and regeneration in sorghum. These candidates span several key functional categories, including AP2/EREBP, GRF, DOF, HB/WOX, ARF, ARR, epigenetic/cell-cycle regulators, and explant-specific factors, and were prioritized by hub connectivity across multiple modules, conservation with functionally characterized orthologs in maize, rice, and *Arabidopsis*, and consistent enrichment in DR-induced samples across both genotypes and explant types (**Table S5**). These candidates were grouped into three priority tiers (Tier 1–3) based on hub connectivity across multiple network modules and/or genotypes, and the strength of supporting ortholog evidence in Arabidopsis, rice, and maize (**see Table S5 legend for full tier criteria**). Because of its recalcitrance, we selected Tx2783 for functional validation. Four genes (*SbHB41, SbEREB157, SbEREB167*, and *SbEREB174*), previously uncharacterized for their role in SE and regeneration in sorghum, were evaluated, with WUS-mediated (pPHP96021 or pPHP96564) embryogenesis as a positive control (**Figure 7**). In the absence of a new DR (control), explants rapidly underwent necrosis following transformation and failed to produce regenerated plants (**Figure 7D**).

**Figure 7.**
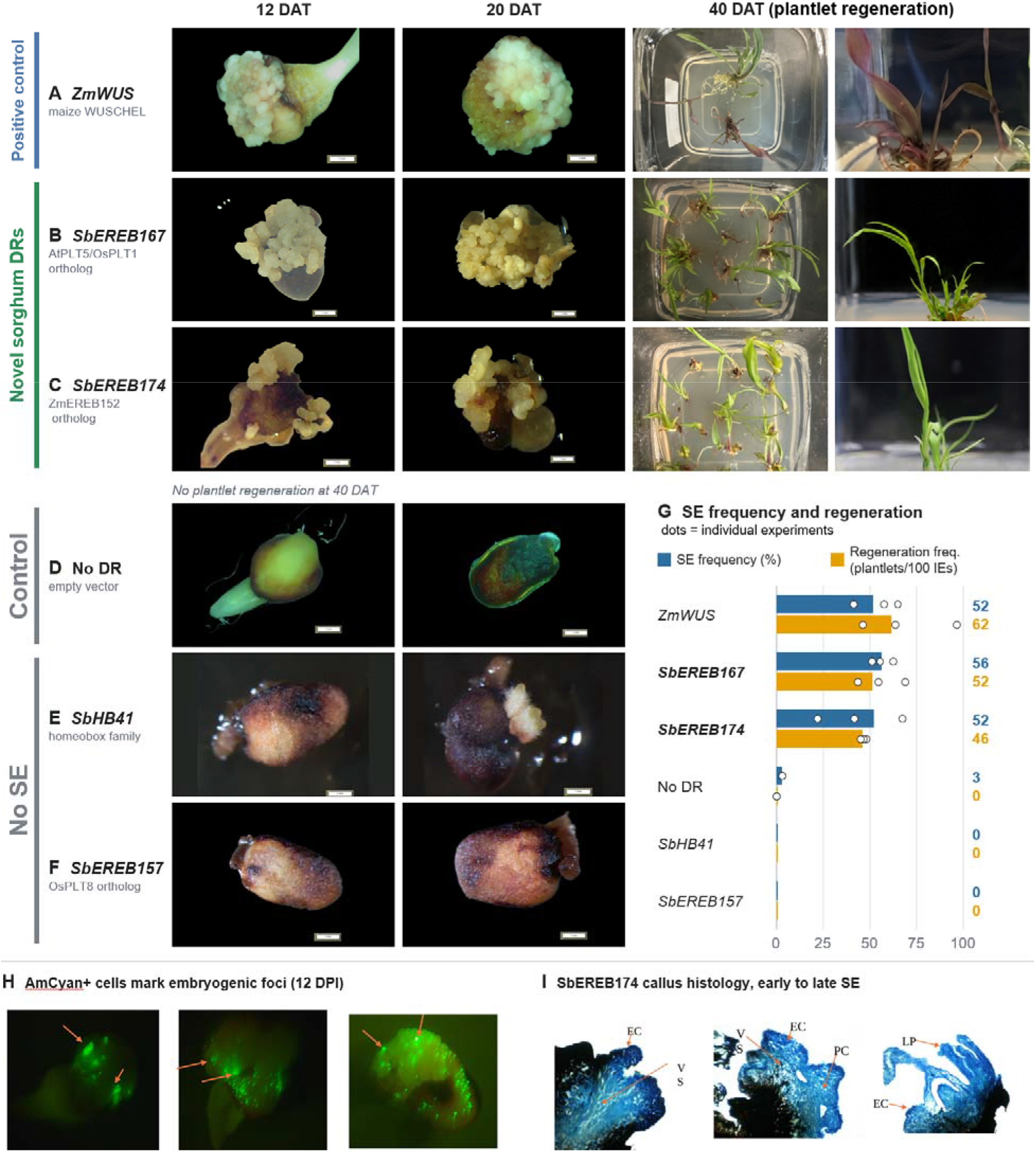
Functional validation of novel sorghum developmental regulators in the recalcitrant genotype Tx2783. **(A–F)** Immature embryos (IEs) transformed with *ZmWUS* (A, positive control), *SbEREB167* (B), *SbEREB174* (C), an empty vector (D, No DR), *SbHB41* (E) or *SbEREB157* (F), imaged at 12 and 20 days after transformation (DAT) and, for A–C, at 40 DAT during plantlet regeneration. Scale bars, 1 mm (12 and 20 DAT). D–F produced no plantlets. **(G)** Somatic embryo (SE) frequency (% of IEs forming SE, blue) and regeneration index (plantlets per 100 IEs, orange), pooled across independent experiments (total events / total IEs; *ZmWUS, SbEREB167 and SbEREB174*, three experiments; No DR, two); dots show individual experiments. *SbHB41* and *SbEREB157* produced no SE or regeneration. Source data are in Table S6. **(H)** *AmCYAN* fluorescence in *ZmWUS*, *SbEREB167* and *SbEREB174* callus at 12 days post infection (DPI); arrows mark fluorescent foci of morphogenic structures (bright-field images in Fig. S19). **(I)** Histology of *SbEREB174* callus at early, intermediate and late embryogenic stages. EC, embryogenic cells; VS, vasculature; PC, parenchymatous callus cells; LP, leaf primordia. SEM images of embryogenic structures are in Fig. S18.

In three independent experiments, *SbEREB174* and *SbEREB167* induced somatic embryogenesis (SE) in 52.3% (138/264) and 56.4% (220/390) of immature embryos, respectively, comparable to the ZmWUS positive control (52.0%, 195/375), whereas empty-vector explants rarely formed SE (3.3%, 6/180; Figure 7A–D, G; Table S6). The three DRs yielded 46.2, 51.5 and 61.9 plantlets per 100 IEs, respectively. AmCYAN fluorescence marked the embryogenic foci (**Figure 7H**), and the induced structures resembled those formed with *WUS* and *WUS+BBM*, including early to late embryogenic stages with vascular tissue and leaf primordia (**Figure 7I; Figures S18, S19**). In contrast, *SbHB41* (Homeobox family) and *SbEREB157* (AP2/EREB family) did not induce visible embryogenic structures, showing a response comparable to control (**Figure 7E-G**). These results indicate that somatic-embryogenesis induction was gene-specific, and not all candidate DRs can induce embryogenic reprogramming. Among the tested genes, we emphasized on *SbEREB167* and *SbEREB174* that promoted rapid SE, with embryogenic structures observed as early as 5 DAT and well-developed embryogenic calli by 12 DAT (**Figure 7**). Overall, two independently selected, previously uncharacterized genes were sufficient to drive robust SE and suggested that our network-based prioritization strategy was a constructive approach for discovering native DR genes in recalcitrant crops. Regenerated T plants grew normally after transfer to soil and showed no obvious pleiotropic phenotypes at this stage, comparable to the *ZmWUS* control **(Figure S21)**. The remaining prioritized candidates in **Table S5** (*GRF, DOF, HB/WOX, ARF/ARR*, and epigenetic/cell-cycle regulators) represent a resource for future functional validation in sorghum and other cereals. Chromatin and cell-cycle regulators (e.g., SbPHD20, SbSNF2.27, SbE2F5/7) and hormone-signaling hubs (SbARF2, SbARR5/9) represent complementary, non-AP2/EREBP entry points that may act synergistically with the validated regulators. Collectively, we identified and characterized novel sorghum genes that improve SE and genetic transformation in a recalcitrant genotype.

## DISCUSSION

This work establishes an effective transformation framework for two previously recalcitrant sorghum genotypes using developmental regulators, significantly expanding the genetic accessibility of sorghum improvement. Additionally, omics analysis supports the hypothesis that sorghum reflects a distinct, defense/stress-dominated transcriptional state rather than a simple quantitative shortfall in regenerative capacity. *WUS, BBM* and *GRF–GIF* actively suppress this competing state while engaging fate-specification and proliferative programs, and the more recalcitrant genotype (Tx2783) required broader and more sustained transcriptional reprogramming. In summary: (1) early timepoints (7 DAT) are dominated by stress and cell-wall remodeling, while later stages (15 DAT) shift toward developmental signaling and regeneration competence; (2) WUS drives fate specification, whereas GRF-GIF preferentially modulates growth, hormone metabolism, and structural remodeling; and (3) both DRs accelerate the transition from a stress-dominated to a developmental transcriptional state. However, it remains unclear whether these differences are driven primarily by genetic variation, the epigenetic landscape, hormone responsiveness, cell wall properties, or interactions among these factors.

Future studies integrating comparative genomics, genome-wide association analyses using diverse sorghum association panels, and chromatin accessibility profiling will be instrumental in identifying the causal genetic and regulatory factors that determine transformability across sorghum germplasm. Although *SbEREB167* and *SbEREB174* efficiently promote somatic embryogenesis and regeneration, their direct downstream targets and molecular mode of action remain determined.

### Transcriptomics insights reveal mechanisms underlying DR-mediated regeneration

Ectopic expression of WUS and BBM is sufficient to induce embryogenic competence in somatic tissue by activating fate-specification and stem-cell-identity genes (Lowe *et al*., 2016; McFarland *et al*., 2023). The GRF-GIF module functions through a complementary route that promotes cell proliferation and meristem regeneration (Debernardi *et al*., 2020; Lee *et al*., 2009; Li *et al*., 2024). The genotype- and time-dependent suppression of defense/secondary-metabolite pathways by both regulators align with the broader concept that developmental reprogramming. The stress due to culture and *Agrobacterium*-induced defense can block establishment of the embryogenic identity unless actively overridden by strong developmental regulators (Duan *et al*., 2018; Gelvin, 2003; Schmidt *et al*., 2025) (**Figure 8**). The Tx2783 (grain) genotype reflects the more extensive transcriptional reprogramming required to overcome its stronger defense and secondary-metabolite states. By comparison, the less pronounced stress response in Tx2910 (forage) reflects its comparatively higher transformation and regeneration efficiency. Together, our data indicate that somatic-cell reprogramming proceeds through an integrated sequence of signal perception, activation of core cell-cycle and division modules, stress and maturation adaptation, and engagement of embryogenic pathways. Cell-cycle regulators, *CDKs*, and *E2Fs* were elevated in DR-treated samples across both explant types (**File S17**), indicating conserved basal cell-cycle regulation during SE.

**Figure 8:**
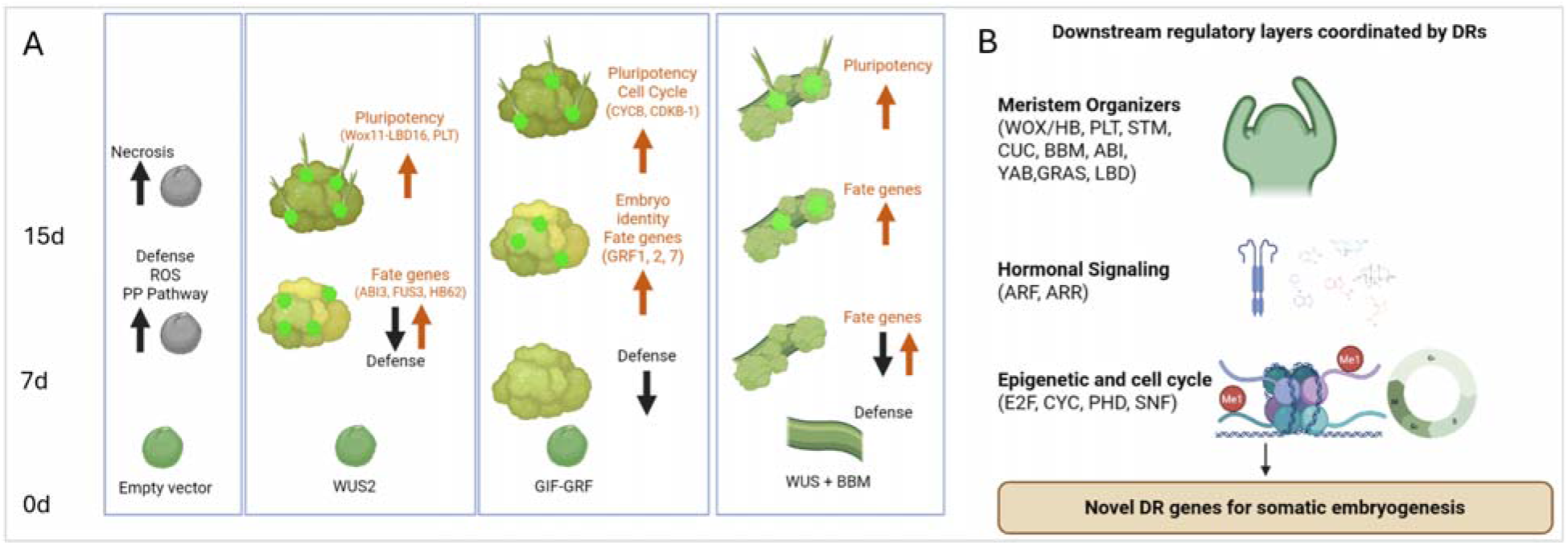
Proposed model of DR-mediated transcriptional reprogramming during SE in sorghum. **(A)** Effect of DR versus no-DR treatment on cellular reprogramming and SE induction. Green dots mark DR/reporter-expressing cells; arrows indicate up-or down-regulation of the indicated gene categories. In Empty vector defense/stress signaling (ROS, PP - phenylpropanoid pathway) remains active and explants progress to necrosis rather than embryogenic callus. In immature embryo explants with WUS2 suppresses defense signaling while activating pluripotency (*WOX11-LBD16*, *PLT*) and fate-specification genes (*ABI3*, *FUS3*, *HB62*) within 7 days of transformation. Immature seed explants with GIF-GRF first suppress defense signaling (7d) and then activate fate genes (*CYCB*, *CDKB-1*) and embryo-identity genes (*GRF1*, *GRF2*, *GRF7*). Leaf explants transformed with WUS+BBM produces synergistic suppression of defense signaling together with activation of pluripotency and fate genes, yielding the robust SE induction. **(B)** Downstream regulatory layers coordinated by DRs, based on GRN/GCN hub-gene analysis showed genes related meristem organizers (*WOX/HB, PLT, STM, CUC, BBM, ABI, YAB, GRAS, LBD*), hormonal signaling (*ARF, ARR*), and epigenetic/cell-cycle regulation (*E2F, CYC, PHD, SNF*). Convergence of these layers prioritized novel DR candidates, including SbEREB167 and SbEREB174 (Figure 7).

snRNA-seq provides cellular-resolution support for these bulk-derived conclusions. The defense-dominant transcriptional state that bulk analysis identified as the default, DR-suppressed baseline in Tx2783 corresponds to a discrete, non-dividing, wound-responsive cell population that DR-treated tissue moves away from, highlighting transitionary states masked by bulk RNA-seq limitations. Likewise, the proliferative module in which we identified *SbEREB167* and *SbEREB174* as bulk network hubs where both genes showed the strongest treatment-induced upregulation. The two datasets therefore converge from complementary angles. Bulk RNA-seq identified genes and pathways change with treatment across the tissue, while snRNA-seq pinpoint that these changes are concentrated in, and plausibly define, the specific proliferative cell population from which regenerated plants likely arise.

Somatic-embryogenesis regulators *ABI3* (*Sobic.007G000400, Sobic.003G398200*), *FUS3* (*Sobic.003G275400*), and *NF-YA1* (*Sobic.001G127800*) were elevated in DR samples in both genotypes (**File S13**). *ABI3*, a B3-family regulator of embryo maturation, promotes embryogenic competence and is characteristically upregulated in dedifferentiated and embryogenic cells (Freitas *et al*., 2019; Kang *et al*., 2021), while *FUS3* cooperates with *LEC* and *ABI3* pathways to maintain embryonic cell identity in *Arabidopsis* (Horstman *et al*., 2017). Similarly, recent work highlights the potential *ABI/FUS2* family genes as positive regulators of regeneration, though equivalent crosstalk in monocots remains poorly characterized (Ramasamy *et al*., 2026). Both genotypes exhibited high expression of transcription factor (TF) families known to regulate plant development and stress responses, including AP2/EREBP, ARF, C2C2-Dof, MYB, NAC, WRKY, MADS-box, bHLH, Homeobox, and GRF, suggesting their involvement in coordinating developmental and embryogenic processes during regeneration (**File S13**). Among the identified hub genes, MYB TFs such as *SbMYB132* (*AtMYB105*) and *SbMYB18* (*AtMYB43*), which regulate cell division in the Arabidopsis shoot meristem (Haga *et al*., 2011; Wang *et al*., 2023a), were detected in both genotypes. AP2/EREBP TFs, including *SbEREB156* (*ZmEREB11*) and *SbEREB49* (*OsERF3*), were identified as hub genes in F_WUS_D15 and in G_WUS_D15/G_WUS_D7, respectively, further supporting their involvement in regeneration. Homeobox/WUS-related TFs, such as *SbHB62* and *SbHB44*, were predominantly active in F_GIF_D15, G_WUS_D7, and G_WUS_D15, consistent with their established roles in maintaining stem-cell identity and promoting embryogenic competence. Additionally, *SbNAC87*, identified as a hub gene in Tx2783 (WUS 7 and 15 DAT), is homologous to *AtNAC030* (VND7), a key regulator of vascular differentiation and xylem formation (Yamaguchi *et al*., 2008) indicating a potential role in vascular tissue development during regeneration. These highly expressed candidate genes along their conserved roles in regeneration demonstrated in dicots species, represent promising regulators for improving sorghum transformation (**File S16**).

### Pleiotropic effects associated with developmental regulators

Constitutive expression of DR can enhance transformation efficiency but may also cause undesirable developmental phenotypes. In wheat, the same *GRF4–GIF1* chimera used here caused only a modest (∼23.9%) reduction in grain number per spike, showing normal fertility (Debernardi *et al*., 2020). In later study, Kronos showed the greatest sensitivity, while Paragon and Cadenza were comparatively less affected in terms of fertility and spikelet number indicating a genotype dependent response to GRF4-GIF over expression (Hayta *et al*., 2025). Similarly, in maize, reduced fertility was associated with *TaGRF4–GIF1/ZmGRF1–GIF1* under the *ZmUbi* promoter (Vandeputte *et al*., 2024), whereas the same chimera expressed under the *Arabidopsis ubiquitin10* promoter did not affect fertility in sorghum cv. Tx430 (Li *et al*., 2024). Consistent with these reports, we observed a significant, consistent reduction in seed number per plant across all independent GRF–GIF events in both Tx2910 and Tx2783 under the strong *ZmUbi* promoter, despite an otherwise normal overall plant phenotype (**Figure S5A, B**). These observations collectively indicate that *GRF–GIF* pleiotropy is strongly influenced by genetic background, promoter choice, and transgene dosage, and support the use of controlled strategies such as tissue-specific promoters (e.g., *ZmPLTP*) or Cre-loxP excision to limit DR expression once its developmental function has been achieved (Hayta *et al*., 2025).

### Potential of Novel DRs for overcoming genotype dependence

DR genes have substantially improved transformation and regeneration in several recalcitrant crops. However, as discussed earlier, their effectiveness varies by genotype, explant type, and the molecular state of the target tissue. Our transcriptomic analysis demonstrates that regeneration competence requires coordinated activation of several regulatory networks at once, such as cell-cycle progression, embryogenic identity, hormone signaling, and suppression of stress responses. Supporting this idea, combinatorial deployment of *PLT5* with *GRF–GIF* recently enabled transformation of several previously recalcitrant plant species (Huo *et al*., 2026), showing that rationally designed DR combinations can extend genotype-independence further than any single regulator alone. Beyond *WUS* and *GRF–GIF*, our integrative GRN/GCN analysis identified a broader set of candidate regulators. Notably, the AP2/EREBP family was disproportionately represented among high-priority, high-connectivity hubs across both genotypes and multiple treatment modules (**Table S5**). This pattern supported our functional results, in which we validated both genes (*SbEREB167* and *SbEREB174*) as efficient inducers of SE. Notably, plants expressing *SbEREB167* or *SbEREB174* appeared phenotypically normal at the seedling stage (Figure S21), although fertility and seed set in the T and T generations remain to be evaluated. *SbEREB174* (ortholog of *ZmEREB152/OsPLT2*) and *SbEREB167* (ortholog of *AtPLT5/OsPLT1*) each integrate auxin–cytokinin signaling and PLT-like pluripotency functions in embryogenic tissue (Renema *et al*., 2025). This is consistent with their strong induction in *WUS* and *GRF–GIF* modules, and with their phylogenetic placement alongside ZmBBM/AtBBM/AtPLT4. *SbEREB167* clustered closely with *ZmBBM* and *SbEREB174* grouped within the broader AtPLT-related clade (**Figure S20A, B**). Because BBM and related PLT proteins are well-established regulators of SE and regeneration, this phylogenetic placement further supports a potential role for these genes in promoting regeneration competence. Despite their shared placement, *SbEREB167* and *SbEREB174* shared only a 65% and 66.1% amino acid identity with *ZmBBM*, respectively. This divergence indicates that *SbEREB167* and *SbEREB174* are not simply orthologs of *ZmBBM*, but distinct, sorghum-regulators, that achieve a similar developmental outcome through non-identical sequences, underscoring their value as novel transformation tools (**Figure S20 B, C**).

## CONCLUSIONS

This study provides a transcriptomic dissection of DR-mediated SE and regeneration in sorghum, revealing that the molecular barrier to transformation in recalcitrant genotypes is not simply a quantitative deficit in regenerative capacity, but a qualitatively distinct transcriptional state. The state is one dominated by persistent stress and must be actively overridden before embryogenic identity can be established. Several mechanistic observations emerged from this work with broad implications for monocot transformation. First, DRs do not operate through a single conserved transcriptional logic but deploy genotype-tailored strategies to reach a common developmental endpoint. In Tx2910, WUS and GRF-GIF acted primarily by suppressing competing stress/defense programs and engaging a relatively accessible embryogenic network, allowing rapid progression through the reprogramming window. In the relatively recalcitrant Tx2783, the same DRs must perform more extensive molecular work, where they actively induced hormone signaling, engaging a broader TF repertoire including DOF, HB, bZIP, and GATA family members, and sustaining a distinct reprogramming transcriptional state. This genotype-tailored mechanism explains why a single DR dosage or timepoint rarely achieves equivalent transformation efficiency across diverse genetic constitutions. Second, the convergence of both genotypes on a conserved late-stage co-expression module (anchored by auxin transport, CLV/WUS-feedback meristem signaling, and CDK-driven cell-cycle progression) defines, a molecular checkpoint that appears to be requirement for sorghum regeneration competence regardless of genotype or explant type. This module represents a tractable target, rather than empirically optimizing entire transformation protocols for each new genotype. Targeted activation of core genes (MEblue4) may offer a more direct, generalizable route to regeneration competence. Third, and most significantly for the broader goal of crop transformation, the integrative GRN analysis identified a set of sorghum-native transcription factors with strong ortholog support and high hub connectivity across multiple independent network analyses. Among several priority genes (**Table S5**), *SbEREB167* and *SbEREB174* (AP2 family) were highly effective and efficient at inducing SE in Tx2783. By identifying and understanding the key barriers, this study lays a mechanistic foundation for developing transformation technologies for precision crop improvement within reach for the full diversity of sorghum and, by extension, other unruly staple crops.

## Data Availability

The raw snRNA-seq sequencing data generated in this study have been deposited in the NCBI Sequence Read Archive (SRA) under bio-project number PRJNA1556803. Analysis scripts used for snRNA-seq processing, clustering, trajectory inference, and co-expression network construction are available from the corresponding author upon reasonable request. The request for LBA4404 Thy- and all pPHP vectors can be made to Nagesh Sardesai. The vector GRF-GIF could be obtained from addgene (https://www.addgene.org/160393/). The request for sorghum specific DRs can be made to Gunvant Patil.

## Author Contributions

PD: methodology, experimental design, formal analysis, data interpretation, and writing – original draft; LD: conducted data analysis, writing, and editing of the manuscript, VD: assisted with sorghum transformation, MJ and LHE: writing, review, and editing of the manuscript; YJ and GP: conceptualization, formal analysis, funding acquisition, supervision, and writing, review and editing.

## Conflict of interest

G.B.P., Y.J, and P.D are co-inventors on a patent application submitted in relation to the findings reported in this study.

## Acknowledgements

We are grateful to the State of Texas’ Governor’s University Research (GURI) for the research funding and partial funding for bulk RNA seq from Sorghum Checkoff. P.D. is grateful to the Texas Tech University Graduate School for Distinguished Graduate Student Assistantship and Texas Tech University Davis College for Helen DeVitt Jones fellowship. The authors thank Dr. Bill Gordon-Kamm and Dr. Nagesh Sardesai (Vylor) for providing PHP vectors and *Agrobacterium* LBA4404-Thy auxotrophic strain for sorghum transformation.

## Supplementary Data

**Figure S1. T-DNA regions of the constructs used in this study.** Schematic maps (not to scale) of the T-DNA of each binary vector (A–K; listed below). Abbreviations: *ZmWUS*, *Zea mays* WUSCHEL; *ZmBBM*, *Z. mays* BABY BOOM; *NptII*, neomycin phosphotransferase II selectable marker; *HptII*, hygromycin phosphotransferase II selectable marker; CmYLCV, Cestrum yellow leaf curling virus promoter; AmCYAN, cyan fluorescent protein from *Anemonia majano*; *rab17*, stress- and ABA-responsive maize promoter; *cre*, Cre recombinase; LTP2, lipid transfer protein 2 promoter; SbBMR, guide RNA targeting *Brown midrib 6*; PINII, proteinase inhibitor II terminator; IN2, maize *In2-1* (*Saf1*) terminator; OCS, octopine synthase terminator; E9, pea *RBCS E9* terminator. RUBY, betalain reporter.

**Figure S2. Tissue sampling scheme for RNA-seq.** Representative explants at each sampling time for IEs and leaf explants transformed with the empty vector (EV), *ZmWUS* + RUBY, *GRF4:GIF1* + AmCyan, or *ZmBBM*/*ZmWUS*. IEs were sampled at 7 and 15 DAT and leaf explants at 7 and 20 DAT (sample codes in Table S1). Images are from Tx2910; Tx2783 was sampled at the same stages.

**Figure S3. RUBY reporter expression in transgenic Tx2910 and Tx2783 plants and T segregation. A–C**, Tx2910 plants expressing *CmYLCV:RUBY* (PHP96021): (A) leaf base/ligule and midrib of mature vegetative plants; (B) spikelets; (C) seeds. Scale bars, 1 cm. **D–F**, as in A–C for Tx2783, showing a similar tissue distribution of betalain pigmentation. **G, H**, T seedlings segregating for RUBY in Tx2910 (G) and Tx2783 (H) events (PHP96021); NTC, non-transgenic control; Ruby-T1, T progeny.

**Figure S4. Genotype-dependent phenolic release in cultured sorghum explants.** Tx2783 IEs (A) and leaf explants (B) release phenolics into the co-cultivation medium, whereas Tx2910 IEs (C) and leaf explants (D) do not. Tx2783 also releases phenolics into rooting medium (E), unlike Tx2910 (F).

**Figure S5. Pleiotropic effects of constitutive DR expression. A, B**, Plants constitutively expressing *ZmUbi:GRF4:GIF1* look phenotypically normal but set few seeds in Tx2783 (A) and none in Tx2910 (B). Scale bar, 1 cm. **C**, Constitutive *ZmBBM*/*ZmWUS* expression (PHP81858) causes thin, curled leaves in Tx2783. **D**, Watery, callus-like structures with deformed roots and shoots

**Figure S6. T_1_ segregation of the AmCyan marker in transgenic sorghum. A, B**, Tx2910 T progeny of PHP81858 events under white light (A) and cyan fluorescence (B). **C, D**, Tx2783 T progeny of PHP81858 events under white light (C) and cyan fluorescence (D). WT, wild-type (non-transformed) control; AmCyan-T1, T progeny of a transgenic event.

**Figure S7. PCR genotyping of randomly selected marker-positive transgenic plants.** PCR with transgene-specific primers on plants selected at random from visible-marker-positive events: *RUBY* in Tx2783 (A) and Tx2910 (B) (PHP96021); *AmCyan* in leaf-derived Tx2783 (C) and Tx2910 (D) plants (PHP81858); and *GIF* in Tx2783 (E) and Tx2910 (F) plants (*GRF4:GIF1*, pPD71). WT, wild-type negative control; P, plasmid positive control; remaining lanes, independent T plants.

**Figure S8. Selection-free transformation mediated by ZmWUS and ZmBBM using the RUBY reporter. A**, T-DNA design of the selection-free vector. **B, C**, RUBY-expressing tissues during embryogenesis and regeneration in Tx2910 (B) and Tx2783 (C). The Tx2783 callus releases abundant phenolics (C). Scale bar, 3 mm.

**Figure S9. Genes upregulated by ZmWUS *in both genotypes.* A**, Venn diagram of genes upregulated relative to the matching no-DR control in *ZmWUS*-expressing Tx2910 and Tx2783 IEs at 7 and 15 DAT, showing genotype-specific and shared DEGs. **B**, Genes upregulated in both genotypes at 7 DAT (387 genes, including *SbYAB7* and *SbEREB174*). **C**, Gene Ontology (GO) biological-process enrichment of the 387 shared genes. **D**, Genes upregulated in both genotypes at 15 DAT. **E**, GO biological-process enrichment of the genes shared at 15 DAT.

**Figure S10. Genes upregulated by GRF4–GIF1 in both genotypes.** Panels A–E as in Fig. S9 for *GRF4–GIF1*-expressing Tx2910 and Tx2783 IEs: (A) Venn diagram of genes upregulated relative to the matching no-DR control at 7 and 15 DAT; (B) genes shared between genotypes at 7 DAT; (C) GO biological-process enrichment of the genes in B; (D) genes shared at 15 DAT; (E) GO enrichment of the genes in D.

**Figure S11. Expression patterns of transcription factor (TF) families during somatic embryogenesis.** Mean Z-scaled expression of TF families across the RNA-seq samples of Tx2910 (A) and Tx2783 (B), with heatmaps of the top 10 members of selected families. *SbEREB174* (OsPLT2 ortholog) and *SbEREB167* (OsPLT1 ortholog) are labelled.

**Figure S12. K-means transcriptional modules responding to ZmBBM + ZmWUS in forage sorghum (Tx2910) leaf explants.** Transcriptional modules identified by k-means clustering in forage sorghum (Tx2910) leaf explants. **A**, Heatmaps of DEGs (log2 fold change ≥ 2; Benjamini–Hochberg-adjusted P < 0.05) clustered into five clusters across the control at 7 DAT (FL_C_D7) and *ZmBBM* + *ZmWUS* at 7 and 20 DAT (FL_BW_D7, FL_BW_D20). **B**, Representative enriched GO biological-process terms for each cluster. **C**, Representative hub TFs for each cluster.

**Figure S13. K-means transcriptional modules responding to ZmBBM + ZmWUS in grain sorghum (Tx2783) leaf explants.** Transcriptional modules identified by k-means clustering in grain sorghum (Tx2783) leaf explants. **A**, Heatmaps of DEGs (log2 fold change ≥ 2; Benjamini–Hochberg-adjusted P < 0.05) clustered into six clusters across the control at 7 DAT (GL_C_D7) and *ZmBBM* + *ZmWUS* at 7 and 20 DAT (GL_BW_D7, GL_BW_D20). **B**, Representative enriched GO biological-process terms for each cluster. **C**, Representative hub TFs for each cluster.

**Figure S14. Weighted gene co-expression network analysis (WGCNA) of leaf explants. A**, Module–trait correlation heatmap for the 18 co-expression modules and the six leaf samples (FL_C_D7, FL_BW_D7, FL_BW_D20, GL_C_D7, GL_BW_D7, GL_BW_D20); the modules fall into three categories of sample-associated expression programs. **B**, Number of genes in each module. **C–F**, GO biological-process enrichment of the modules associated with FL_BW_D7 (C), FL_BW_D20 (D), GL_BW_D7 (E) and GL_BW_D20 (F).

**Figure S15. Single-nucleus RNA-seq of EV and ZmBBM + ZmWUS (BW) tissue. A**, Fraction of nuclei from EV and BW samples in each of the 21 clusters. **B**, UMAP of the integrated dataset split by sample, coloured by cluster; the red circle marks a BW-specific cluster. **C**, Feature plots of transgene expression (*ZmBBM* and *AmCyan*) split by sample; darker colour indicates higher expression. **D**, GO biological-process terms enriched among cluster marker genes (FDR < 0.05). **E**, Feature plots of cell-cycle phase scores (G1/G0, S, G2/M); darker colour indicates higher scores.

**Figure S16. Expression of soybean cell-type marker orthologs in the sorghum snRNA-seq dataset.** UMAP feature plots showing expression of marker genes for cell types defined in published soybean seed and shoot apical meristem (SAM) snRNA-seq datasets: early-maturity (EM), cotyledonary-stage (CS) and middle-maturity (MM) seeds, and SAM. Source datasets:doi:10.1016/j.cell.2024.10.050 and bioRxiv, doi:10.1101/2024.07.08.602332. Marker genes are listed in Supplementary Data.

**Figure S17. Expression of rice cell-type marker orthologs in the sorghum snRNA-seq dataset.** UMAP feature plots showing expression of marker genes for rice cell types: mesophyll, phloem, epidermis, callus founder cell/scutellar epithelium, initial cells, proliferating cells and SAM niche. Source dataset: doi:10.1186/s12870-026-08596-6. Marker genes are listed in Supplementary Data.

**Figure S18. Morphology and histology of somatic embryogenesis induced by DRs. A**, Tx2783 IEs expressing *ZmWUS*, *ZmWUS* + *ZmBBM*, *SbEREB174* or *SbEREB167* at 7 DAT, with scanning electron microscopy (SEM) images of the corresponding embryogenic structures. Scale bar, 1 mm. **B–D**, Histological sections of *SbEREB174*-overexpressing calli at early (B), intermediate (C) and late (D) embryogenic stages. EC, embryogenic cells; PC, parenchymatous callus cells; VS, vasculature; LP, leaf primordia.

**Figure S19. Early detection of embryogenic foci with the AmCYAN marker.** Tx2783 IEs expressing *ZmWUS* + AmCYAN (PHP96564), *SbEREB174* + AmCYAN or *SbEREB167* + AmCYAN at 12 DAT. AmCYAN fluorescence marks transgenic callus and enables early detection of embryo formation; arrows indicate foci of morphogenic structure development.

**Figure S20. Phylogeny and sequence alignment of the 15 candidate regulators. A**, Maximum-likelihood tree of the 15 sorghum candidates (blue) with genes associated with pluripotency in Arabidopsis, rice and maize. **B**, Tree of AP2/EREBP-family genes and related developmental regulators. **C**, Sequence alignment of the sorghum AP2/EREBP candidates and related regulators to the reference sequence (ZmBBM2/ZmEREB187) is indicated. Protein sequences were aligned with MUSCLE and trees were inferred by maximum likelihood in R.

**Figure S21: Morphology of representative T transgenic plants after transfer to soil.** Tx2783 plants regenerated from immature embryos expressing *ZmWUS* (positive control; label “W”), *SbEREB167* (pPD137; label “137”) or *SbEREB174* (pPD136; label “136”). Three independent events per construct are shown. Plants were photographed after hardening in soil (2 weeks after transfer). All plants show normal leaf and shoot morphology without or deformed shoots.

### Supplementary Tables

**Table S1: Sample designations for bulk RNA-seq.** Each sample name encodes the explant and genotype, the developmental regulator (DR) treatment and the sampling time in the format explant/genotype_DR_D[DAT]. Prefixes: F, forage sorghum Tx2910 immature embryos (IEs); G, grain sorghum Tx2783 IEs; FL, Tx2910 leaf explants; GL, Tx2783 leaf explants. DR codes: C, control (empty vector, no DR); GIF, GRF4–GIF1 chimera; WUS, ZmWUS; BW, ZmBBM + ZmWUS. D7, D15 and D20 denote 7, 15 and 20 days after transformation (DAT).

**Table S2: Transformation efficiency, regeneration index, transgenic purity and escape frequency by genotype, explant type and DR treatment:** Immature embryos (IEs) and leaf explants of Tx2910 and Tx2783 were transformed with the constructs indicated (EV, empty vector without a DR; WUS, *ZmWUS*; GRF-GIF, *GRF4–GIF1*; BBM_WUS, *ZmBBM* + *ZmWUS*). Expt, independent experiment. “No. of IEs” is the number of IEs transformed; for leaf explants the denominator is the number of starting seedlings [confirm]. Values are given as number (percentage) where a percentage is shown.

**Table S3: Numbers of differentially expressed genes (DEGs) in pairwise comparisons between sampling times, DR treatments and genotypes.** Each comparison is shown as a pair of sample names (defined in Table S1) with the number of DEGs beneath each name. The number under a sample is the count of genes with significantly higher expression in that sample than in its comparison partner (equivalently, downregulated in the partner). Comparisons cover (i) developmental progression (D15 vs D7), (ii) DR treatment vs control at the same DAT, (iii) genotype (Tx2910 vs Tx2783) and (iv) leaf-explant time courses.

**Table S4. Quality metrics for the snRNA-seq dataset:** Minimum, median, mean and maximum number of unique molecular identifiers (UMIs) and of detected genes per nucleus after quality filtering, for the empty-vector control (EV; no DR) and the ZmBBM + ZmWUS (BW) samples, and for both samples combined. A total of 7,059 nuclei were retained (3,761 EV; 3,298 BW), and 26,599 distinct genes were detected across the combined dataset.

**Table S5: Prioritized candidate regulators of somatic embryogenesis and regeneration:** Candidates were selected from co-expression and gene regulatory network analyses on the basis of hub connectivity, differential regulation across tissues and time points, and functional evidence from orthologs. Genes are grouped by family or functional category. Gene ID, sorghum locus with gene name; Ortholog, closest characterized ortholog in Arabidopsis (At), rice (Os) or maize (Zm); Key modules, RNA-seq sample-specific modules (sample names as in Table S1) or leaf co-expression network modules (named by colour) in which the gene is a hub; Functional relevance, supporting evidence with references. Priority tiers. Tier 1, hub in ≥ 3 modules and/or in both genotypes, with strong ortholog evidence. Tier 2, significant hub in ≥ 2 modules, supported by ortholog function. Tier 3, explant-or genotype-specific, with weaker cross-system support. SbEREB174, SbEREB167, SbEREB157 and SbHB41 were tested functionally in this study (Fig. 7; Table S6); the remaining candidates have yet to be validated by transformation.

**Table S6: Somatic embryogenesis and plant regeneration from Tx2783 immature embryos expressing novel DRs.** Tx2783 IEs were transformed with ZmWUS (positive control), SbEREB167, SbEREB174 or no DR (negative control). Each row is one independent experiment (Expt). No. of IEs, explants transformed. Somatic embryo frequency (%) = (IEs forming somatic embryos ÷ IEs transformed) × 100, scored from AmCyan-positive embryogenic structures (marked “AmCyan positive”) or, where AmCyan was not used, from embryogenic calli identified by morphology (marked “Embryogenic calli”) at [DAT]. Total regenerated plants = number of plantlets recovered per experiment. The regeneration index plotted in Fig. 7G is the pooled number of plantlets per 100 IEs (total plantlets/total IEs x 100). “No regeneration”: no plantlets were recovered. SbHB41 and SbEREB157 produced no somatic embryos or plants.

