## Supplementary Tables for "Developmental regulator-mediated transcriptional reprogramming reveals novel regulators of somatic embryogenesis in sorghum"

**Table S1: Sample designations for bulk RNA-seq.** Each sample name encodes the explant and genotype, the developmental regulator (DR) treatment and the sampling time in the format [explant/genotype]_[DR]_D[DAT]. Prefixes: F, forage sorghum Tx2910 immature embryos (IEs); G, grain sorghum Tx2783 IEs; FL, Tx2910 leaf explants; GL, Tx2783 leaf explants. DR codes: C, control (empty vector, no DR); GIF, GRF4–GIF1 chimera; WUS, ZmWUS; BW, ZmBBM + ZmWUS. D7, D15 and D20 denote 7, 15 and 20 days after transformation (DAT).

| **Sample Name** | **Genotype** | **Explant** | **Developmental Regulator** | **DAT** |
| --- | --- | --- | --- | --- |
| F_C_D7 | Tx2910 | Immature Embryo | None | 7 |
| F_C_D15 | Tx2910 | Immature Embryo | None | 15 |
| F_GIF_D7 | Tx2910 | Immature Embryo | GRF-GIF | 7 |
| F_GIF_D15 | Tx2910 | Immature Embryo | GRF-GIF | 15 |
| F_WUS_D7 | Tx2910 | Immature Embryo | WUS | 7 |
| F_WUS_D15 | Tx2910 | Immature Embryo | WUS | 15 |
| G_C_D7 | Tx2783 | Immature Embryo | None | 7 |
| G_C_D15 | Tx2783 | Immature Embryo | None | 15 |
| G_GIF_D7 | Tx2783 | Immature Embryo | GRF-GIF | 7 |
| G_GIF_D15 | Tx2783 | Immature Embryo | GRF-GIF | 15 |
| G_WUS_D7 | Tx2783 | Immature Embryo | WUS | 7 |
| G_WUS_D15 | Tx2783 | Immature Embryo | WUS | 15 |
| FL_C_D7 | Tx2910 | Leaf | None | 7 |
| FL_BW_D7 | Tx2910 | Leaf | BBM + WUS | 7 |
| FL_BW_D20 | Tx2910 | Leaf | BBM + WUS | 20 |
| GL_C_D7 | Tx2783 | Leaf | None | 7 |
| GL_BW_D7 | Tx2783 | Leaf | BBM + WUS | 7 |
| GL_BW_D20 | Tx2783 | Leaf | BBM + WUS | 20 |

**Table S2: Transformation efficiency, regeneration index, transgenic purity and escape frequency by genotype, explant type and DR treatment:** Immature embryos (IEs) and leaf explants of Tx2910 and Tx2783 were transformed with the constructs indicated (EV, empty vector without a DR; WUS, *ZmWUS*; GRF-GIF, *GRF4–GIF1*; BBM_WUS, *ZmBBM* + *ZmWUS*). Expt, independent experiment. “No. of IEs” is the number of IEs transformed; for leaf explants the denominator is the number of starting seedlings [confirm]. Values are given as number (percentage) where a percentage is shown.

| **Genotype/Explant/DR** | **No. of IEs** | **Transformation efficiency (%)** | **Total Regenerated plants**  **(Regeneration index)** | **Transformation frequency or Transgenic purity (%)** | **Escapes** |
| --- | --- | --- | --- | --- | --- |
| Tx2910/IE/EV (Expt1) | 100 | 7 | No regeneration |  |  |
| Tx2783/IE/EV (Expt1) | 100 | 5 | No regeneration |  |  |
| Tx2783/IE/EV (Expt2) | 50 | 2 | No regeneration |  |  |
| Tx2910/IE/WUS (Expt 1) | 40 | 28 (70%) | 108 (2.7) | 86 (79.62%) | 22 |
| Tx2783/IE/WUS (Expt 1) | 50 | 13 (26%) | 32 (0.64) | 28 (87.5%) | 4 |
| Tx2783/IE/WUS (Expt 2) | 50 | 15 (30%) | 36 (0.72) | 29 (80.55%) | 7 |
| Tx2783/IE/WUS (Expt 3) | 100 | 46 (46%) | 120 (1.2) | 112 (93.33%) | 8 |
| Tx2783/IE/WUS (Total from above 3 independent experiments) | 200 | 74 (37%) | 188 (0.94) | 169 (89.89) | 19 |
| Tx2910/IE/GRF-GIF (Expt 1) | 75 | 38 (50.66%) | 110 (1.46) | 84 (76.36%) | 26 |
| Tx2783/IE/GRF-GIF (Expt 1) | 65 | 32 (49.23%) | 84 (1.26) | 68 (80.95%) | 16 |
| Genotype/Explant/DR | No. of starting seedlings | Transformation efficiency (%) | Total Regenerated plants (Regeneration index) | **Transformation frequency** or Transgenic purity (%) | Escapes |
| Tx2910/Leaf/BBM_WUS (Expt 1) | 18 | 9 (50%) | 41 (2.27) | 31 (75.6%) | 10 |
| Tx2783/Leaf/BBM_WUS (Expt 1) | 25 | 11 (44%) | 38 (1.52) | 37 (97.36%) | 1 |

**Transformation efficiency (%)** = (Number of IEs producing embryogenic structures with visible marker gene expression 7 DAT/ Total number of starting explants) × 100

**Regeneration Index=** Total isolated plantlets per No. of starting seedlings reported as a unitless ratio (plantlets per explant) rather than a percentage; because a single embryogenic callus can give rise to more than one shoot, this index is expected to routinely exceed 1 and should not be expressed as a bounded percentage

**Transformation frequency or Transgenic purity (%)** = (Number of Ruby⁺ or AmCyan⁺ plantlets with visible expression as well as PCR confirmation for transgene/Total number of regenerated explants) × 100

**Escapes** refer to regenerated plantlets lacking visible marker expression as well as PCR confirmation for transgene.

IE- Immature embryo

EV- Empty vector (Without DR genes)

**Table S3: Numbers of differentially expressed genes (DEGs) in pairwise comparisons between sampling times, DR treatments and genotypes.** Each comparison is shown as a pair of sample names (defined in Table S1) with the number of DEGs beneath each name. The number under a sample is the count of genes with significantly higher expression in that sample than in its comparison partner (equivalently, downregulated in the partner). Comparisons cover (i) developmental progression (D15 vs D7), (ii) DR treatment vs control at the same DAT, (iii) genotype (Tx2910 vs Tx2783) and (iv) leaf-explant time courses.

| Sample pairs | |
| --- | --- |
| F_C_D15 | **F_C_D7** |
| 1907 | 1155 |
| F_GIF_D15 | **F_C_D15** |
| 3718 | 2580 |
| F_WUS_D15 | **F_C_D15** |
| 1163 | 2527 |
| F_C_D15 | **G_C_D15** |
| 8045 | 5068 |
| F_GIF_D7 | **F_C_D7** |
| 482 | 507 |
| F_WUS_D7 | **F_C_D7** |
| 808 | 1360 |
| G_C_D7 | **F_C_D7** |
| 2225 | 2636 |
| G_C_D15 | **G_GIF_D15** |
| 3661 | 6596 |
| G_C_D15 | **G_Wus_D15** |
| 3625 | 6912 |
| G_C_D7 | **G_GIF_D7** |
| 99 | 161 |
| G_C_D7 | **G_Wus_D7** |
| 2120 | 1920 |
| FL_BW_D7 | **FL_C_D7** |
| 8151 | 6322 |
| GL_BW_D20 | **GL_BW_D7** |
| 3330 | 4991 |
| FL_BW_D7 | **FL_BW_D20** |
| 8481 | 6535 |
| GL_BW_D7 | **GL_C_D7** |
| 4895 | 3476 |

**Table S4. Quality metrics for the snRNA-seq dataset:** Minimum, median, mean and maximum number of unique molecular identifiers (UMIs) and of detected genes per nucleus after quality filtering, for the empty-vector control (EV; no DR) and the ZmBBM + ZmWUS (BW) samples, and for both samples combined. A total of 7,059 nuclei were retained (3,761 EV; 3,298 BW), and 26,599 distinct genes were detected across the combined dataset.

| Sample |  |  |  |  |
| --- | --- | --- | --- | --- |
| UMIs | **Min.** | **Median** | **Mean** | **Max.** |
| EV | 563 | 1047 | 1576.4 | 17295 |
| BM | 426 | 671.5 | 873.5 | 7743 |
| Unique Genes | **Min.** | **Median** | **Mean** | **Max.** |
| EV | 448 | 723 | 871.7 | 3897 |
| BM | 343 | 514 | 607.1 | 2553 |
| combined | **Min.** | **Median** | **Mean** | **Max.** |
| UMIs | 426 | 839 | 1248.1 | 17295 |
| Unique Genes | 343 | 606 | 748.1 | 3897 |
|  | **nuclei** |  |  |  |
| EV | 3761 |  |  |  |
| BM | 3298 |  |  |  |
| Gene Coverage | 26599 |  |  |  |

**Table S5: Prioritized candidate regulators of somatic embryogenesis and regeneration:** Candidates were selected from co-expression and gene regulatory network analyses on the basis of hub connectivity, differential regulation across tissues and time points, and functional evidence from orthologs. Genes are grouped by family or functional category. Gene ID, sorghum locus with gene name; Ortholog, closest characterized ortholog in Arabidopsis (At), rice (Os) or maize (Zm); Key modules, RNA-seq sample-specific modules (sample names as in Table S1) or leaf co-expression network modules (named by colour) in which the gene is a hub; Functional relevance, supporting evidence with references. Priority tiers. Tier 1, hub in ≥ 3 modules and/or in both genotypes, with strong ortholog evidence. Tier 2, significant hub in ≥ 2 modules, supported by ortholog function. Tier 3, explant- or genotype-specific, with weaker cross-system support. SbEREB174, SbEREB167, SbEREB157 and SbHB41 were tested functionally in this study (Fig. 7; Table S6); the remaining candidates have yet to be validated by transformation.

| Gene ID | Ortholog | Priority | Key modules | Functional relevance |
| --- | --- | --- | --- | --- |
| AP2/EREBP — embryogenic cell proliferation & axis establishment | | | | |
| Sobic.010G215100  (SbEREB174) | ZmEREB15  OsPLT2 | Tier 1 | F-C-7, F-C-15, F-GIF-7, G-GIF-15, G-WUS-7 | Auxin–cytokinin signaling in embryogenic tissue; Hub in BBM/WUS callus (Renema *et al.*, 2025) |
| Sobic.003G390600  (SbEREB165) | ZmBBM  AtAP2 | Tier 1 | G_GIF_D15, G_GIF_D7, G_WUS_D15 | Sorghum BBM homolog; hub across grain DR modules |
| Sobic.002G022600 (SbEREB161)  Sobic.001G075700 (SbEREB157) | OsPLT8 | Tier 1 | WUS and GIF modules, both genotypes | Conserved PLT-like roles; embryogenic proliferation (Maqbool *et al.*, 2025) |
| Sobic.001G036800  (SbEREB156) | ZmEREB11 | Tier 2 | G_WUS_D7, F_WUS_D15 | Control of inflorescence and floral meristem establishment (Chuck *et al.*, 2008) |
| Sobic.006G245500  (SbEREB167) | OsPLT1  AtPLT5 | Tier 2 | WUS module, both genotypes | Maintain PIN1 expression at meristem periphery; somatic embryogenesis (Prasad *et al.*, 2011; Tsuwamoto *et al.*, 2010) |
| GRFs — Growth Regulatory factors | | | | |
| Sobic.006G203400  (SbGRF7) | ZmGRF5 | Tier 1 | F_WUS_D7, F_WUS_D15, G_C_D15, G_GIF_D15 | Homolog of ZmGRF5; hub across 4 modules spanning both genotypes and timepoints |
| Sobic.001G139800  (SbGRF2)  Sobic.004G269900  (SbGRF4) | AtGRF2/4 | Tier 2 | F_GIF_D7, F_WUS_D7/ F_GIF_D7, F_WUS_D7, G_GIF_15 | Hubs in forage early-stage; promotes meristematic competence (Lee *et al.*, 2026; Rodriguez *et al.*, 2016) |
| DOF — vascular differentiation & developmental plasticity | | | | |
| Sobic.004G266200  (SbDOF19) | OsDOF7  AtDOF5.6 | Tier 1 | F_C_D7, G_WUS_D7, F_GIF_D15 | Recurrent DOF hub across IE and leaf explant networks in both genotypes; conserved in wheat somatic embryogenesis (Liu *et al.*, 2023) |
| Sobic.006G267900  (SbDOF23) | OsDOF17  AtDOF5.4 | Tier 2 | FL_C_D7, GL_C_D7, GL_BW_D7; | Enriched in both genotype control and BW networks; broad developmental plasticity role (Liu *et al.*, 2023) |
| Sobic.008G001700  (SbDOF25) | OsDOF12  AtDOF1.5 | Tier 2 | G_GIF_D15, G_GIF_D7, G_WUS_D15 | Regulates vascular cell differentiation and lignin biosynthesis (Ramachandran *et al.*, 2020) |
| HB/WOX — stem cell maintenance & shoot meristem reprogramming | | | | |
| Sobic.006G254900  (SbHB62) | AtWUS | Tier 1 | G_WUS_D7, G_WUS_D15, F_GIF_D15 | WUS ortholog; candidate for ectopic embryogenic identity induction (Xu *et al.*, 2005) |
| Sobic.007G087600  (SbHB65) | OsWOX10 | Tier 1 | F_GIF_D7/D15, G_GIF_D7/D15 | Broad regeneration competence role. Known role in root primordia development (Kawai *et al.*, 2022). |
| Sobic.010G030200  (SbHB78) | ZmHB77  AtHAT4 | Tier 2 | G_C_D7, G_GIF_D15, F_GIF_D15, F_GIF_D7 | Modulates auxin distribution and tissue patterning during embryo formation (Turchi *et al.*, 2013). |
| Sobic.003G336600  (SbHB41) | OsWOX8  AtWOX13 | Tier 2 | F_WUS_D7 | Early embryogenic identity marker (Ogura *et al.*, 2023) |
| Sobic.003G360200  (SbHB44) | OsWOX9  AtWOX7 | Tier 1 | F_GIF_D15, G_WUS_D7, G_WUS_D15 | Promote embryogenic competence (Kamiya *et al.*, 2003; Yang *et al.*, 2024) |
| ARF/ARR — auxin–cytokinin signaling axis | | | | |
| Sobic.010G073600  (SbARF2) | AtARF7 | Tier 1 | F_GIF_D7, G_GIF_D7, G_WUS_D7, G_WUS_D15 | Bridges auxin–cytokinin networks; regulates LBD genes (Okushima *et al.*, 2007) |
| Sobic.010G064700  (SbARR9)  Sobic.004G330900  (SbARR5) | AtARR12 | Tier 2 | F_GIF_D7/D15, G_GIF_D7 | Type-B response regulators integrating cytokinin signaling into embryogenic activation. AtARR12 activates WUS expression (Dai *et al.*, 2017; Ishida *et al.*, 2008) |
| Epigenetic & cell-cycle — mitotic maintenance & chromatin remodeling | | | | |
| Sobic.004G243900  (SbE2F5)  Sobic.006G008800  (SbE2F7) | AtE2F | Tier 3 | G_GIF_D7 | Maintain mitotic activity (Vlieghe *et al.*, 2005) |
| Sobic.003G439700  (SbPHD20)  Sobic.004G009600  (SbSNF2.27) | AtRX5  AtSNF2 | Tier 2 | G_WUS_D7/F_C_D15, G_C_D15, G_GIF_D15, G_GIF_D7 | Cell-cycle regulation; epigenetic regulation of boundary formation; CUC gene activation (Luo *et al.*, 2024; Raynaud *et al.*, 2006) |
| Explant-specific & conserved grass regulators | | | | |
| Sobic.001G041500  (SbbHLH1) | ZmbHLH48 | Tier 1 | F_GIF_D7, G_WUS_D7, G_GIF_D15 | Regulatory framework for meristem identity (Renema *et al.*, 2025) |
| Sobic.002G224200  (SbHSF9) | AtSCZ | Tier 1 | blue4 leaf GCN; GL_BW_D20 | Stem-cell division and cell patterning (Pernas *et al.*, 2010) |
| Sobic.007G165100  (SbLBD27) | AtLBD29 | Tier 3 | blue leaf GCN modules | Regulators of differentiation and cell-fate transition (Feng *et al.*, 2012) |
| Meristem organization and organ patterning | | | | |
| Sobic.010G003700  (SbGRAS73) | AtSCL6/OsHAM4 | Tier 2 | F_C_D15, F_GIF_D7, G_WUS_D15, G_C_D7, | Maintenance of shoot apical meristem indeterminacy (Engstrom *et al.*, 2011; Fan *et al.*, 2015) |
| Sobic.003G008700  (SbMYB132) | AtMYBR-1 | Tier 2 | G_WUS_D7, F_WUS_D7 | Controls the number and size of the stem cell pools at the shoot apex and root tip (Wang *et al.*, 2023) |
| Sobic.006G160800  (SbYAB7) | OsYAB5/AtYAB1 | Tier 1 | F_C_D7/D15, F_WUS_D7/D15, G_GIF_D7, G_WUS_D15 | Non-cell autonomous meristem factor (Goldshmidt *et al.*, 2008) |
| Sobic.007G000400  (SbABI41) | OsIDEF1  AtLEC2 | Tier 3 | G_GIF_D15 | Induces the formation of somatic embryos and other organ-like structures (Stone *et al.*, 2008) |

**Table S6: Somatic embryogenesis and plant regeneration from Tx2783 immature embryos expressing novel DRs.** Tx2783 IEs were transformed with ZmWUS (positive control), SbEREB167, SbEREB174 or no DR (negative control). Each row is one independent experiment (Expt). No. of IEs, explants transformed. Somatic embryo frequency (%) = (IEs forming somatic embryos ÷ IEs transformed) × 100, scored from AmCyan-positive embryogenic structures (marked “AmCyan positive”) or, where AmCyan was not used, from embryogenic calli identified by morphology (marked “Embryogenic calli”) at [DAT]. Total regenerated plants = number of plantlets recovered per experiment. The regeneration index plotted in Fig. 7G is the pooled number of plantlets per 100 IEs (total plantlets ÷ total IEs × 100). “No regeneration”: no plantlets were recovered. SbHB41 and SbEREB157 produced no somatic embryos or plants.

| DR gene | No. of IEs | Somatic embryo (frequency %) | Total Regenerated plants |
| --- | --- | --- | --- |
| No DR (Expt. 1) | 120 | 4 (3.33 %) | No regeneration |
| No DR (REP 2) | 60 | 2 (3.33%) | No regeneration |
| ZmWUS (Expt.1) | 155 | 64 (*AmCyan* positive) 41.29% | 72 |
| ZmWUS (Expt 2) | 60 | 39 (*AmCyan* positive) 65% | 58 |
| ZmWus (Expt 3) | 160 | 92 (*AmCyan* positive) 57.5% | 102 |
| SbEREB167 (Expt. 1) | 180 | 92 (Embryogenic calli) 51.11% | 98 |
| SbEREB167 (Expt 2) | 45 | 25 (Embryogenic calli) 55.5% | 31 |
| SbEREB167 (Expt 3) | 165 | 103 (*AmCyan* positive) 62.42% | 72 |
| SbEREB174 (Expt. 1) | 54 | 12 (Embryogenic calli) 22.22% | 26 |
| SbEREB174 (Expt 2) | 60 | 25 (Embryogenic calli) 41.66% | 28 |
| SbEREB174 (Expt 3) | 150 | 101 (*AmCyan* positive) 67.33% | 68 |
