## Supplementary Figures for "Developmental regulator-mediated transcriptional reprogramming reveals novel regulators of somatic embryogenesis in sorghum"

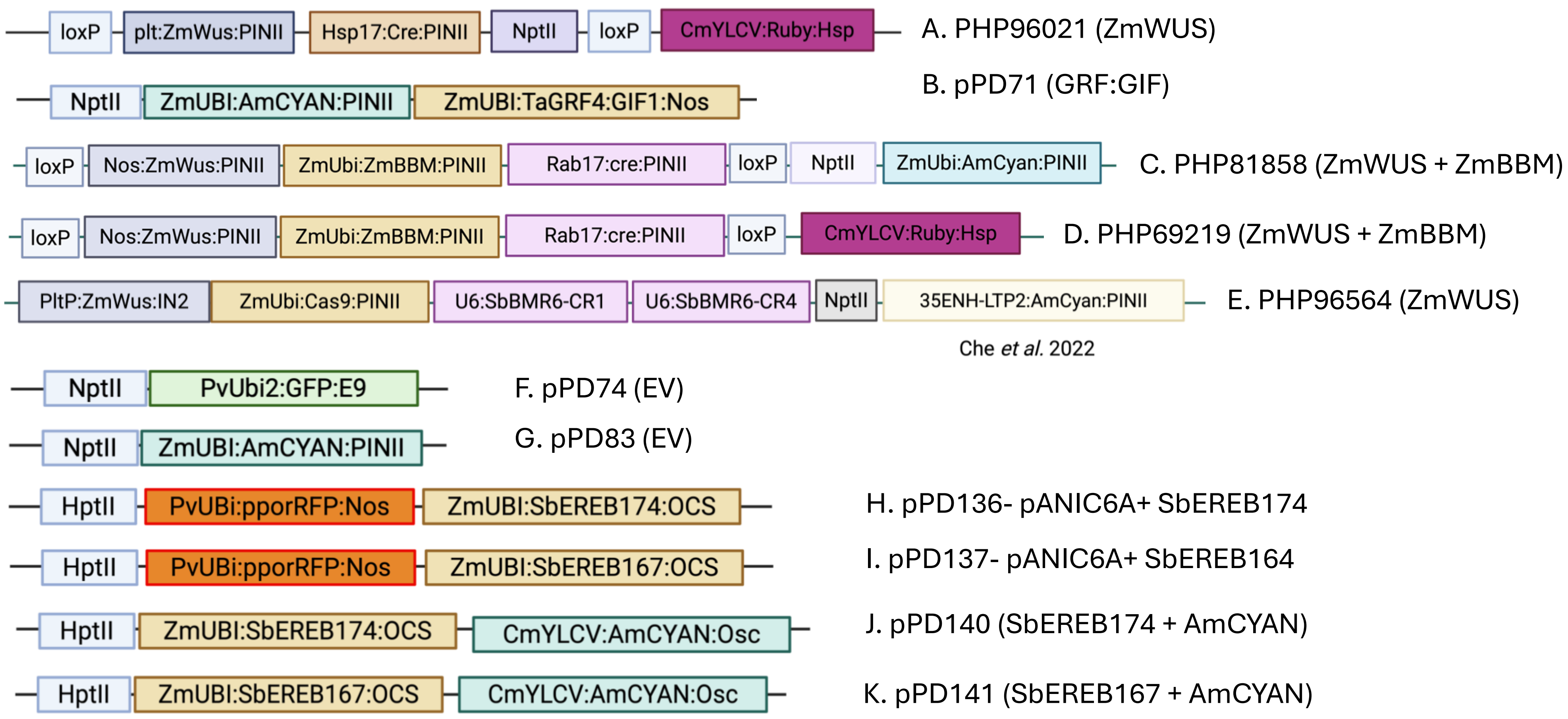


**Figure S1. T-DNA regions of the constructs used in this study.** Schematic maps (not to scale) of the T-DNA of each binary vector (A–K; listed below). Abbreviations: *ZmWUS*, *Zea mays* WUSCHEL; *ZmBBM*, *Z. mays* BABY BOOM; *NptII*, neomycin phosphotransferase II selectable marker; *HptII*, hygromycin phosphotransferase II selectable marker; CmYLCV, Cestrum yellow leaf curling virus promoter; AmCYAN, cyan fluorescent protein from *Anemonia majano*; *rab17*, stress- and ABA-responsive maize promoter; *cre*, Cre recombinase; LTP2, lipid transfer protein 2 promoter; SbBMR, guide RNA targeting *Brown midrib 6*; PINII, proteinase inhibitor II terminator; IN2, maize *In2-1* (*Saf1*) terminator; OCS, octopine synthase terminator; E9, pea *RBCS E9* terminator. RUBY, betalain reporter.

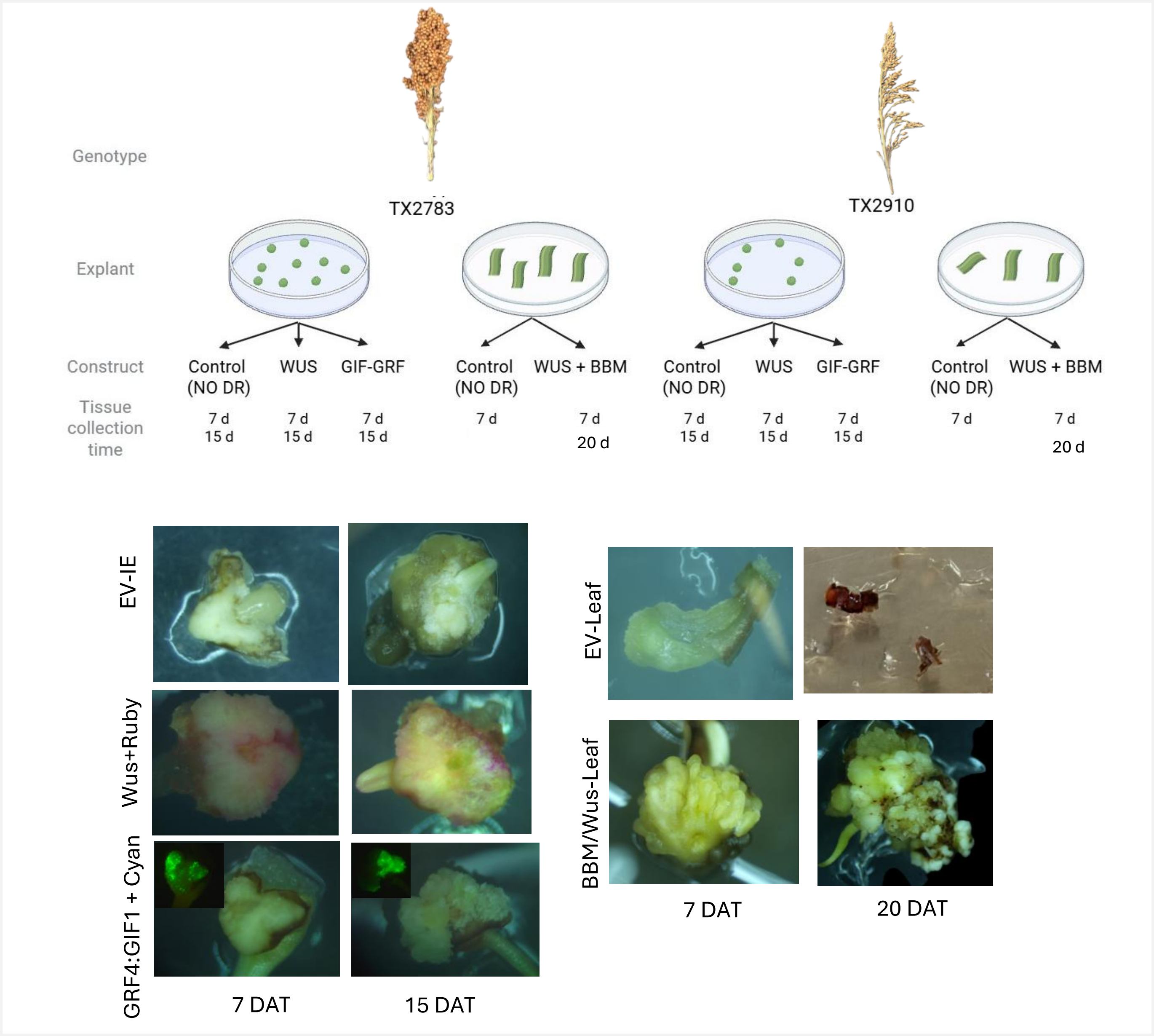


**Figure S2. Tissue sampling scheme for RNA-seq.** Representative explants at each sampling time for IEs and leaf explants transformed with the empty vector (EV), *ZmWUS* + RUBY, *GRF4:GIF1* + AmCyan, or *ZmBBM/ZmWUS*. IEs were sampled at 7 and 15 DAT and leaf explants at 7 and 20 DAT (sample codes in Table S1). Images are from Tx2910; Tx2783 was sampled at the same stages.

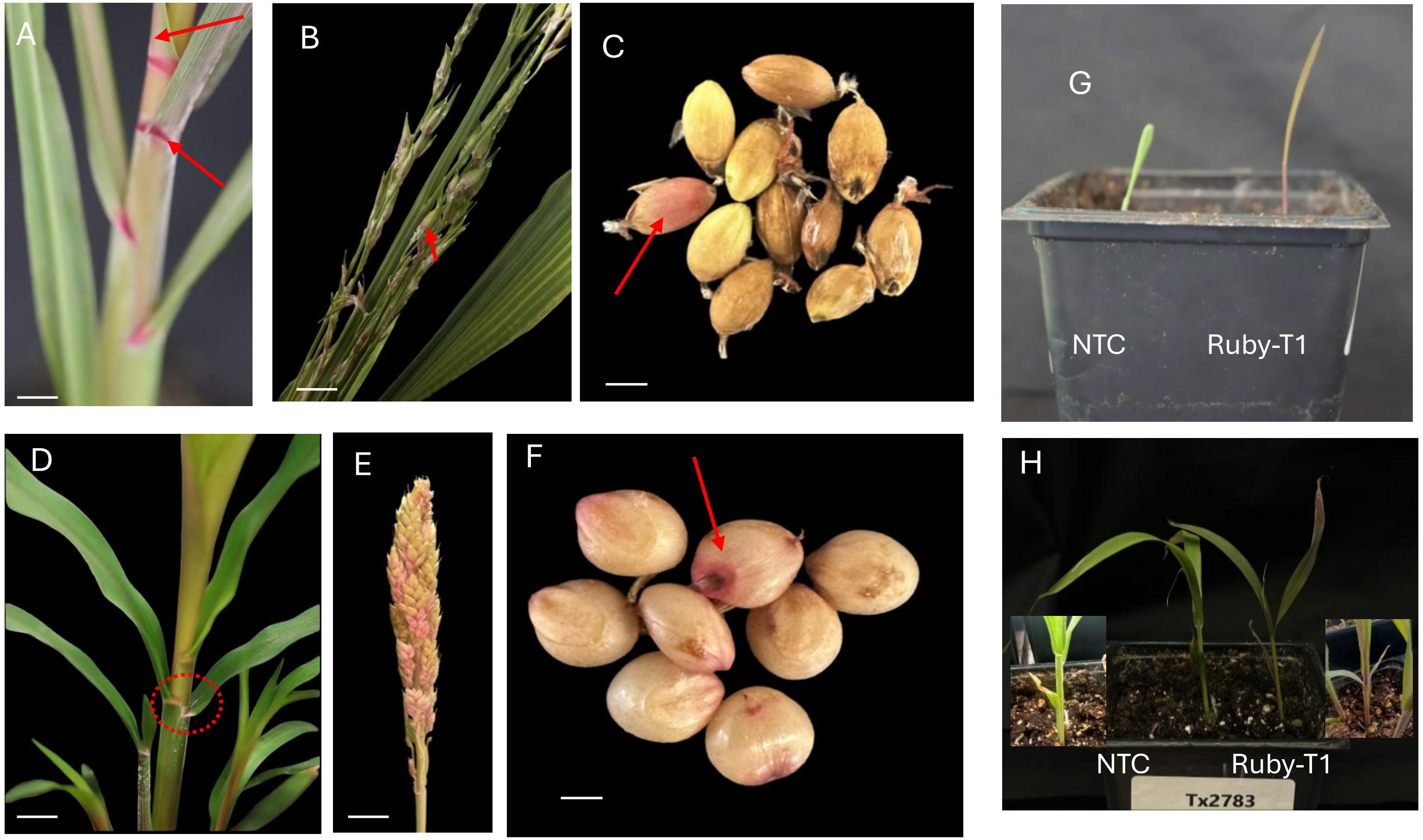


**Figure S3. RUBY reporter expression in transgenic Tx2910 and Tx2783 plants and $T_1$ segregation.**

**A–C**, Tx2910 plants expressing *CmYLCV:RUBY* (PHP96021): (A) leaf base/ligule and midrib of mature vegetative plants; (B) spikelets; (C) seeds. Scale bars, 1 cm. **D–F**, as in A–C for Tx2783, showing a similar tissue distribution of betalain pigmentation. **G, H**, $T_1$ seedlings segregating for RUBY in Tx2910 (G) and Tx2783 (H) events (PHP96021); NTC, non-transgenic control; Ruby-T1, $T_1$ progeny.

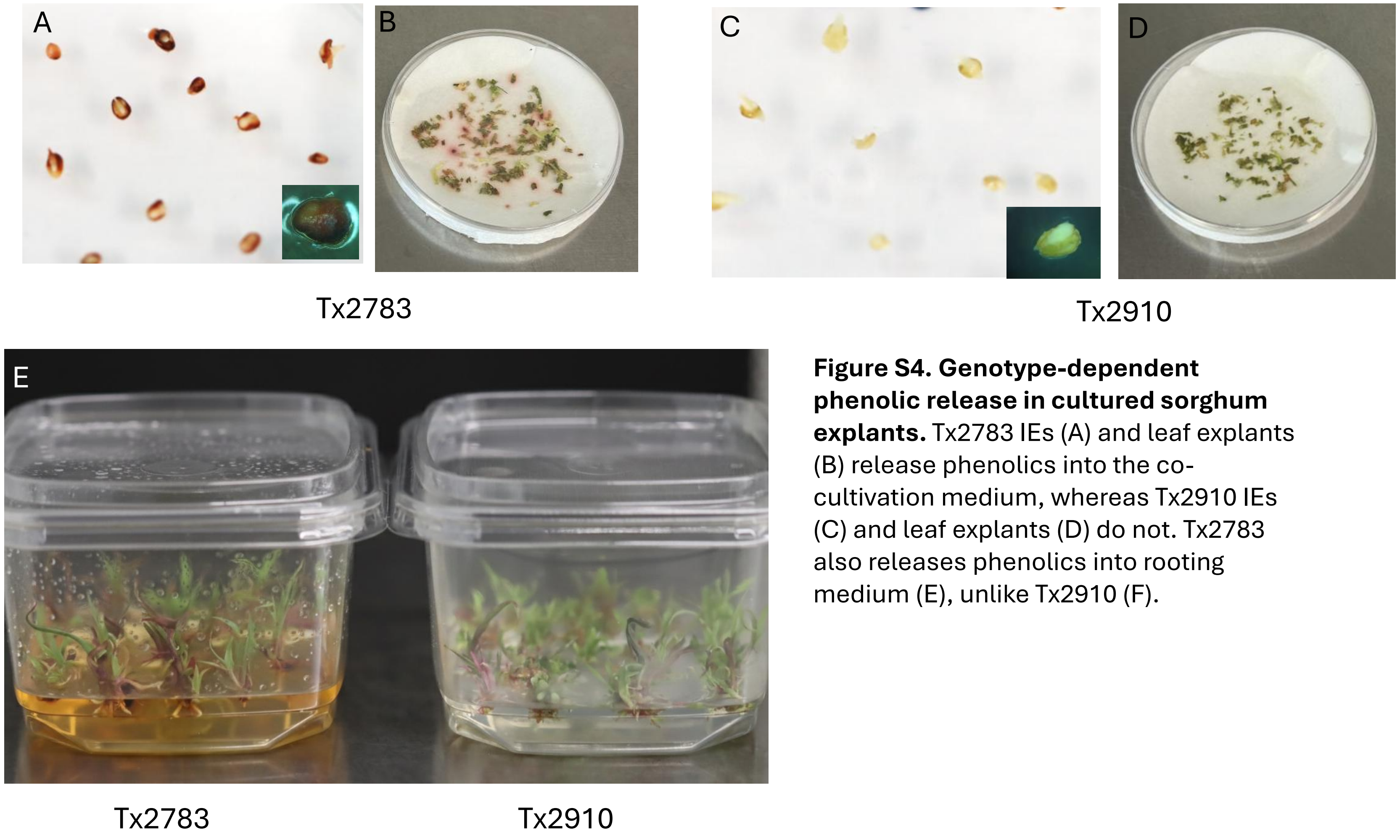


**Figure S4. Genotype-dependent phenolic release in cultured sorghum explants.** Tx2783 IEs (A) and leaf explants (B) release phenolics into the co-cultivation medium, whereas Tx2910 IEs (C) and leaf explants (D) do not. Tx2783 also releases phenolics into rooting medium (E), unlike Tx2910 (F).

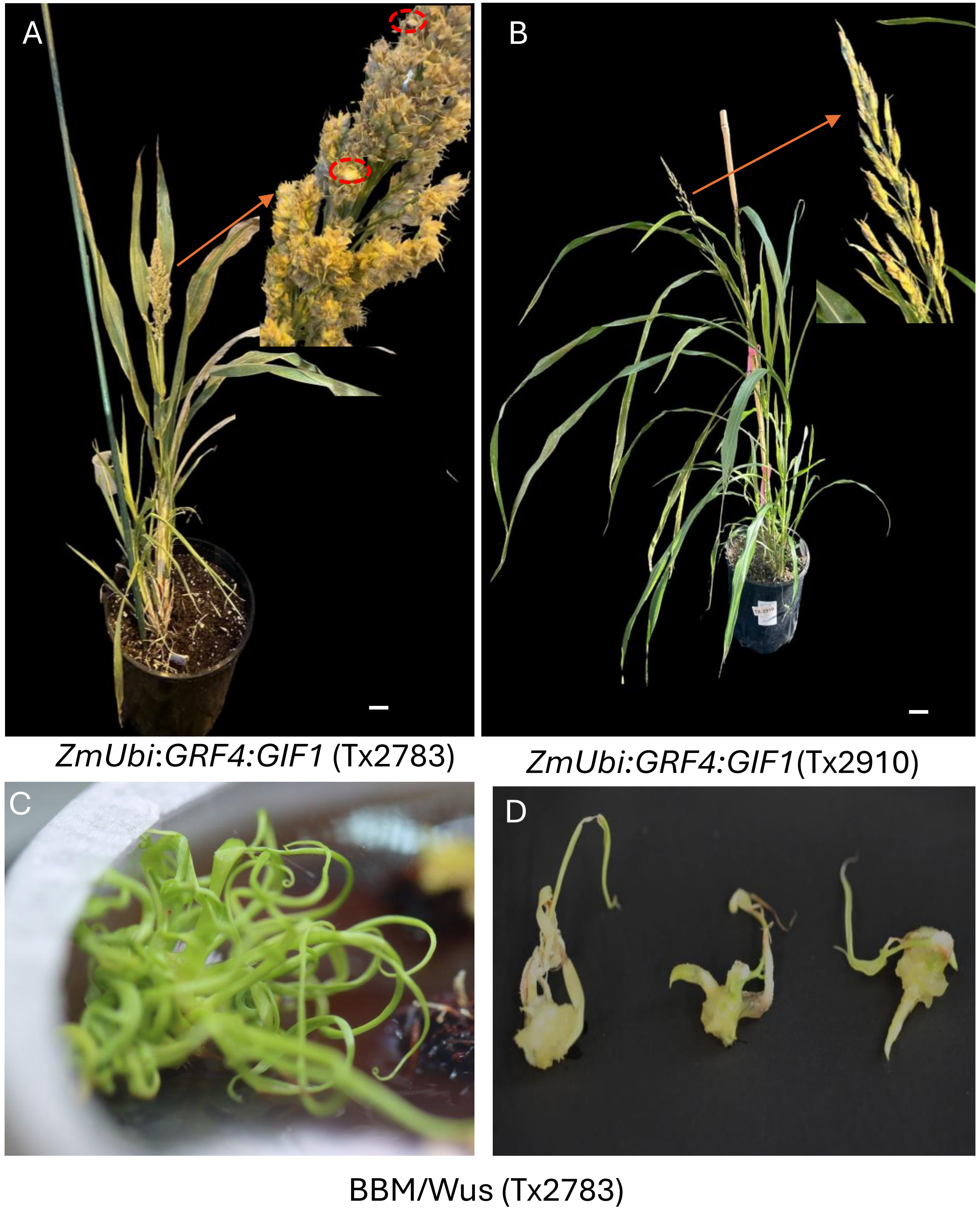


**Figure S5. Pleiotropic effects of constitutive DR expression.**

**A, B**, Plants constitutively expressing *ZmUbi:GRF4:GIF1* look phenotypically normal but set few seeds in Tx2783 (A) and none in Tx2910 (B). Scale bar, 1 cm. **C**, Constitutive *ZmBBM/ZmWUS* expression (PHP81858) causes thin, curled leaves in Tx2783. **D**, Watery, callus-like structures with deformed roots and shoots

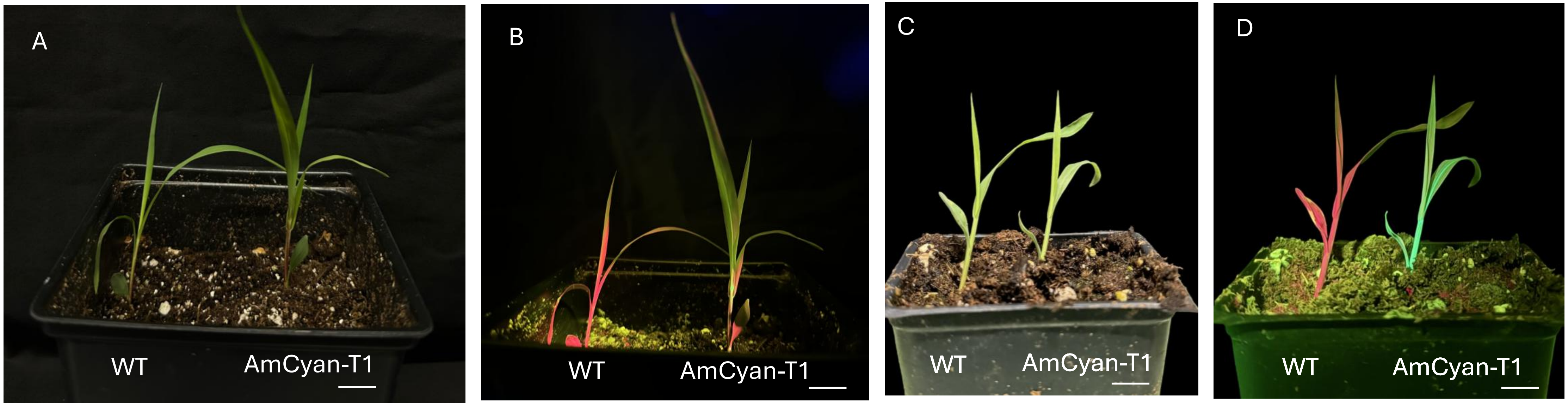


**Figure S6. $T_1$ segregation of the AmCyan marker in transgenic sorghum. A, B**, Tx2910 $T_1$ progeny of PHP81858 events under white light (A) and cyan fluorescence (B). **C, D**, Tx2783 $T_1$ progeny of PHP81858 events under white light (C) and cyan fluorescence (D). WT, wild-type (non-transformed) control; AmCyan-T1, $T_1$ progeny of a transgenic event.

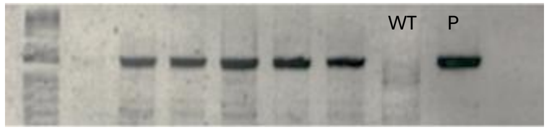


A. Ruby specific PCR for Tx2783 (PHP96021)

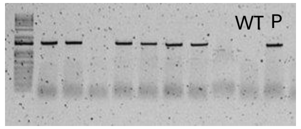


B. Ruby specific PCR for Tx2910 (PHP96021)

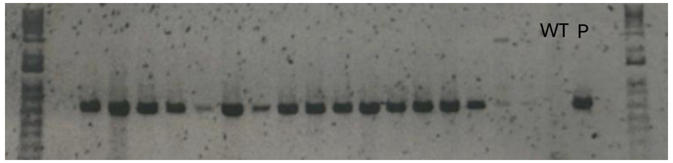


C. *AmCyan* specific PCR for Tx2783 (Leaf, PHP81858)

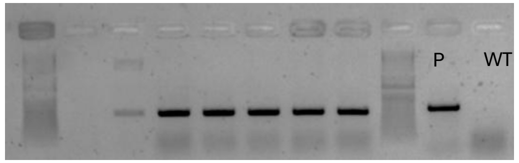


D. *AmCyan* specific PCR for Tx2910 (Leaf, PHP81858)

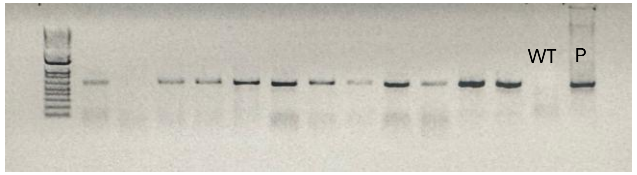


E. *GIF* specific PCR for Tx2783 (GIF:GRF pPD71)

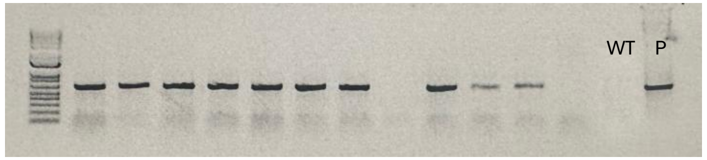


F. *GIF* specific PCR for Tx2910 (GIF:GRF pPD71)

**Figure S7. PCR genotyping of randomly selected marker-positive transgenic plants.** PCR with transgene-specific primers on plants selected at random from visible-marker-positive events: *RUBY* in Tx2783 (A) and Tx2910 (B) (PHP96021); *AmCyan* in leaf-derived Tx2783 (C) and Tx2910 (D) plants (PHP81858); and *GIF* in Tx2783 (E) and Tx2910 (F) plants (*GRF4:GIF1*, pPD71). WT, wild-type negative control; P, plasmid positive control; remaining lanes, independent $T_0$ plants.

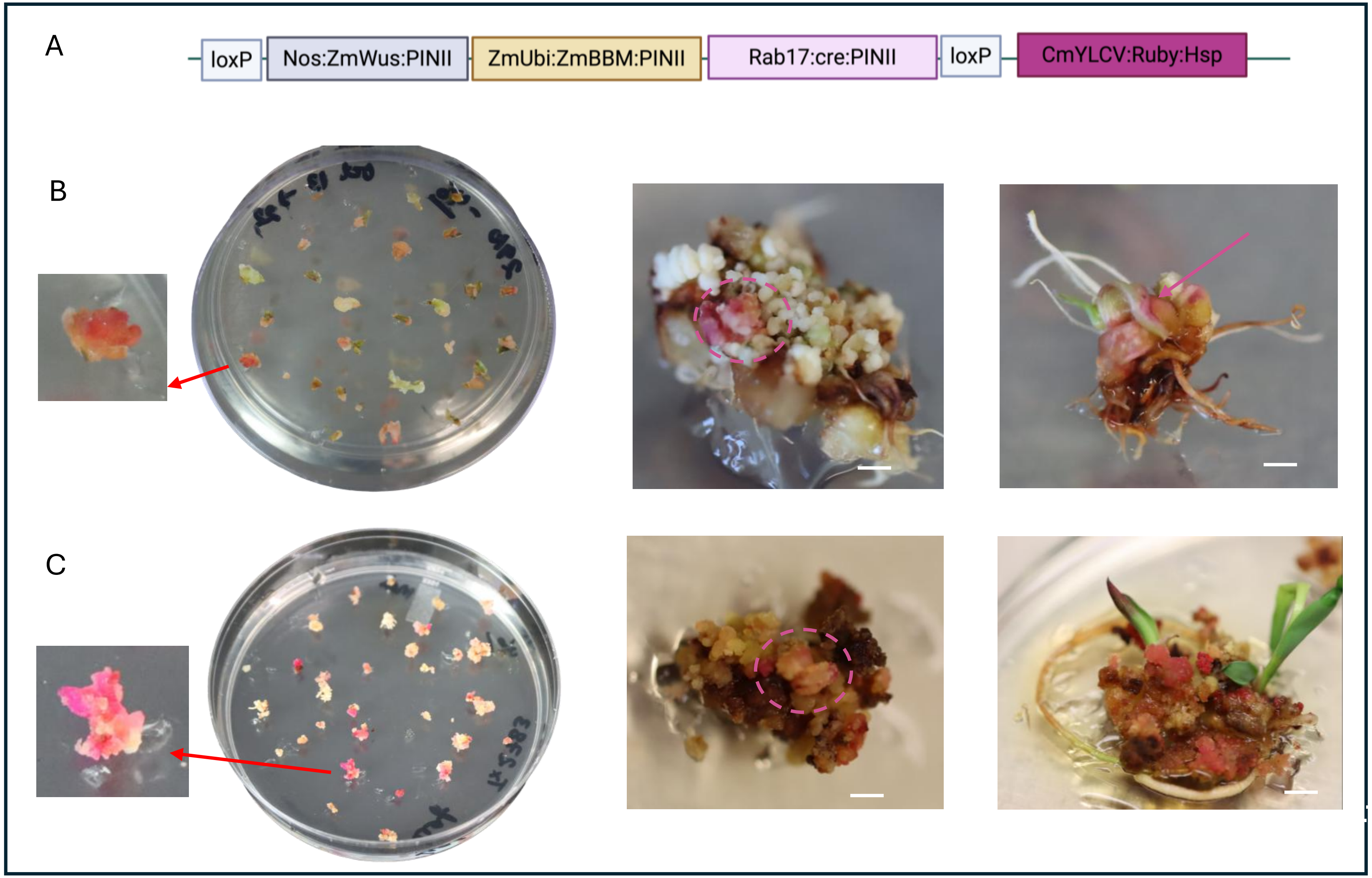


**Figure S8. Selection-free transformation mediated by ZmWUS and ZmBBM using the RUBY reporter. A**, T-DNA design of the selection-free vector. **B, C**, RUBY-expressing tissues during embryogenesis and regeneration in Tx2910 (B) and Tx2783 (C). The Tx2783 callus releases abundant phenolics (C). Scale bar, 3 mm.

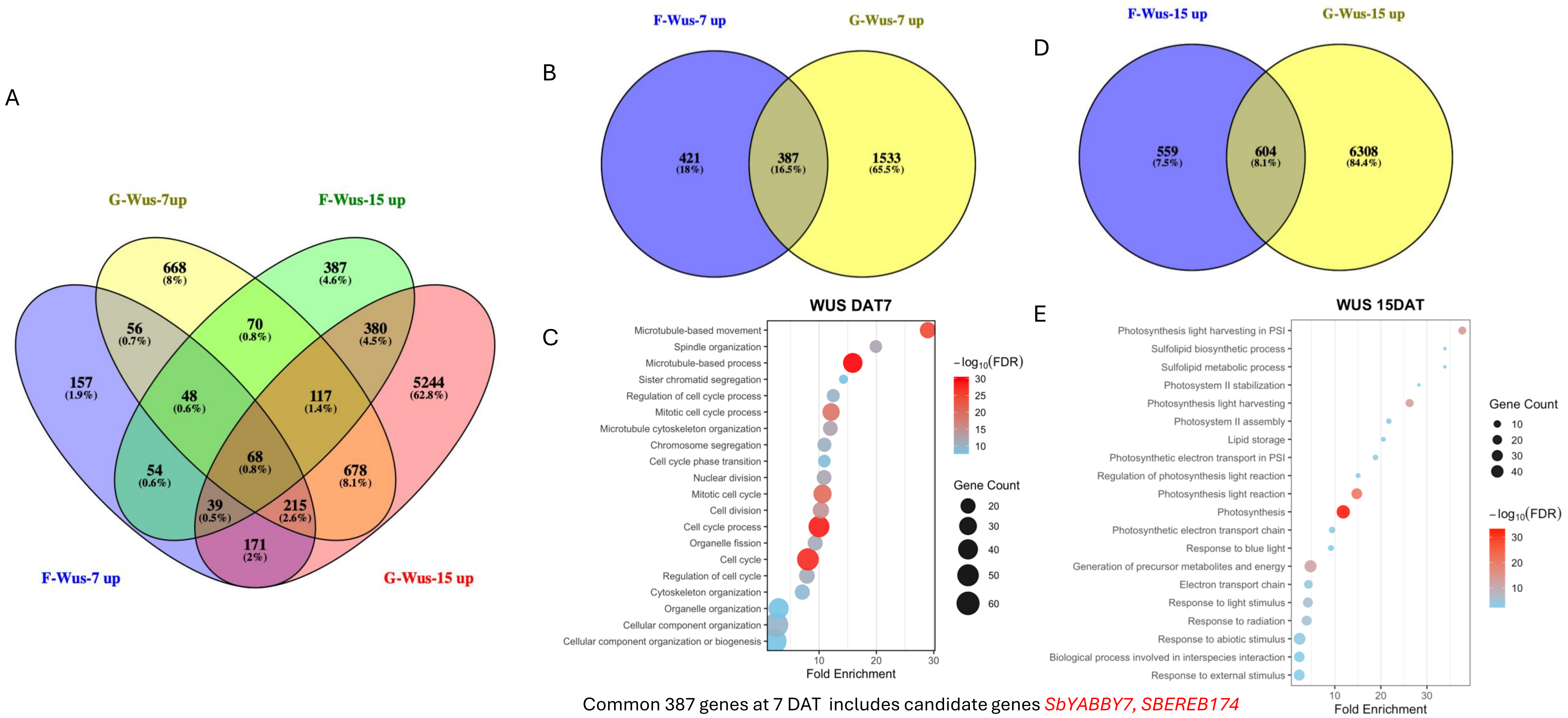


Common 387 genes at 7 DAT includes candidate genes *SbYABBY7, SBEREB174*

**Figure S9. Genes upregulated by ZmWUS *in both genotypes*. A**, Venn diagram of genes upregulated relative to the matching no-DR control in *ZmWUS*-expressing Tx2910 and Tx2783 IEs at 7 and 15 DAT, showing genotype-specific and shared DEGs. **B**, Genes upregulated in both genotypes at 7 DAT (387 genes, including *SbYAB7* and *SbEREB174*). **C**, Gene Ontology (GO) biological-process enrichment of the 387 shared genes. **D**, Genes upregulated in both genotypes at 15 DAT. **E**, GO biological-process enrichment of the genes shared at 15 DAT.

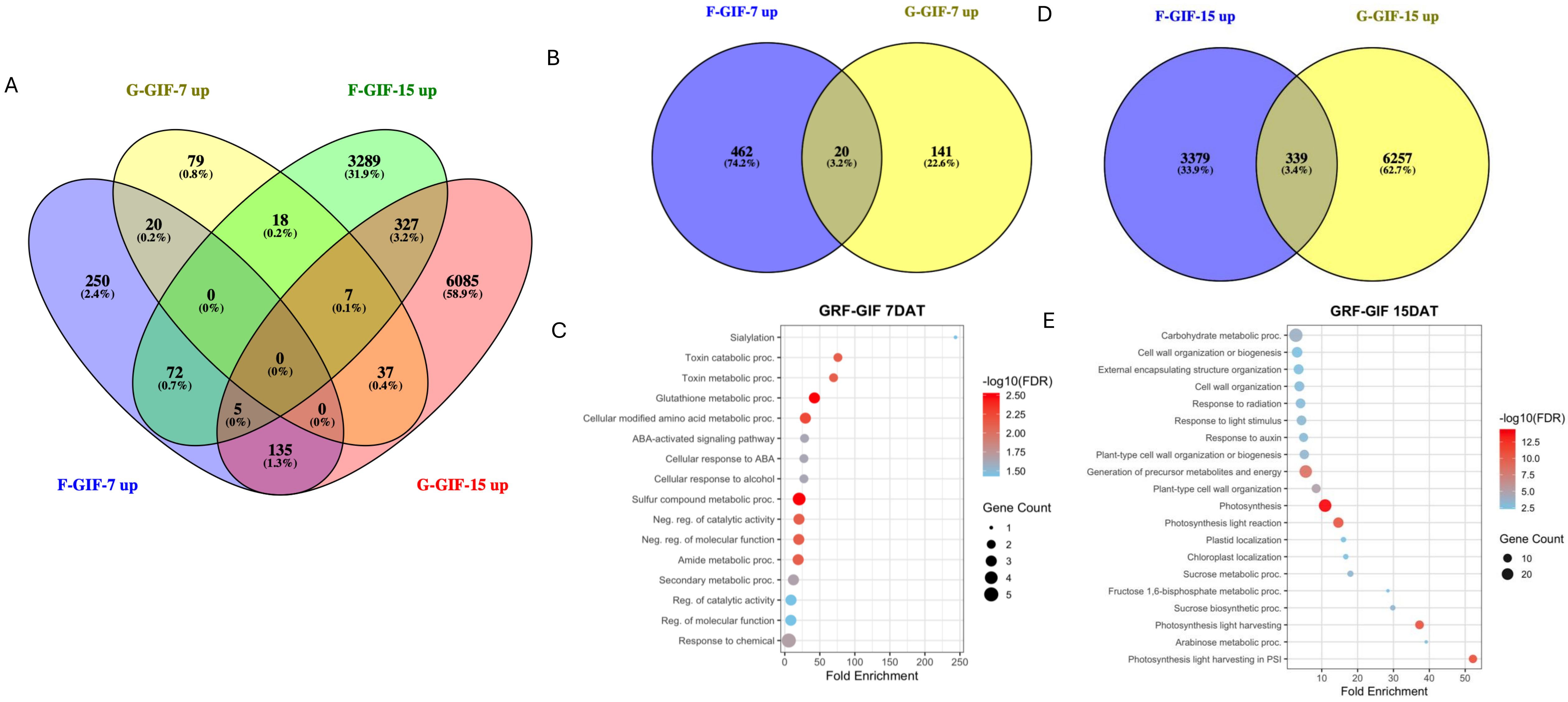


**Figure S10. Genes upregulated by GRF4–GIF1 in both genotypes.** Panels A–E as in Fig. S9 for *GRF4–GIF1*-expressing Tx2910 and Tx2783 IEs: (A) Venn diagram of genes upregulated relative to the matching no-DR control at 7 and 15 DAT; (B) genes shared between genotypes at 7 DAT; (C) GO biological-process enrichment of the genes in B; (D) genes shared at 15 DAT; (E) GO enrichment of the genes in D.

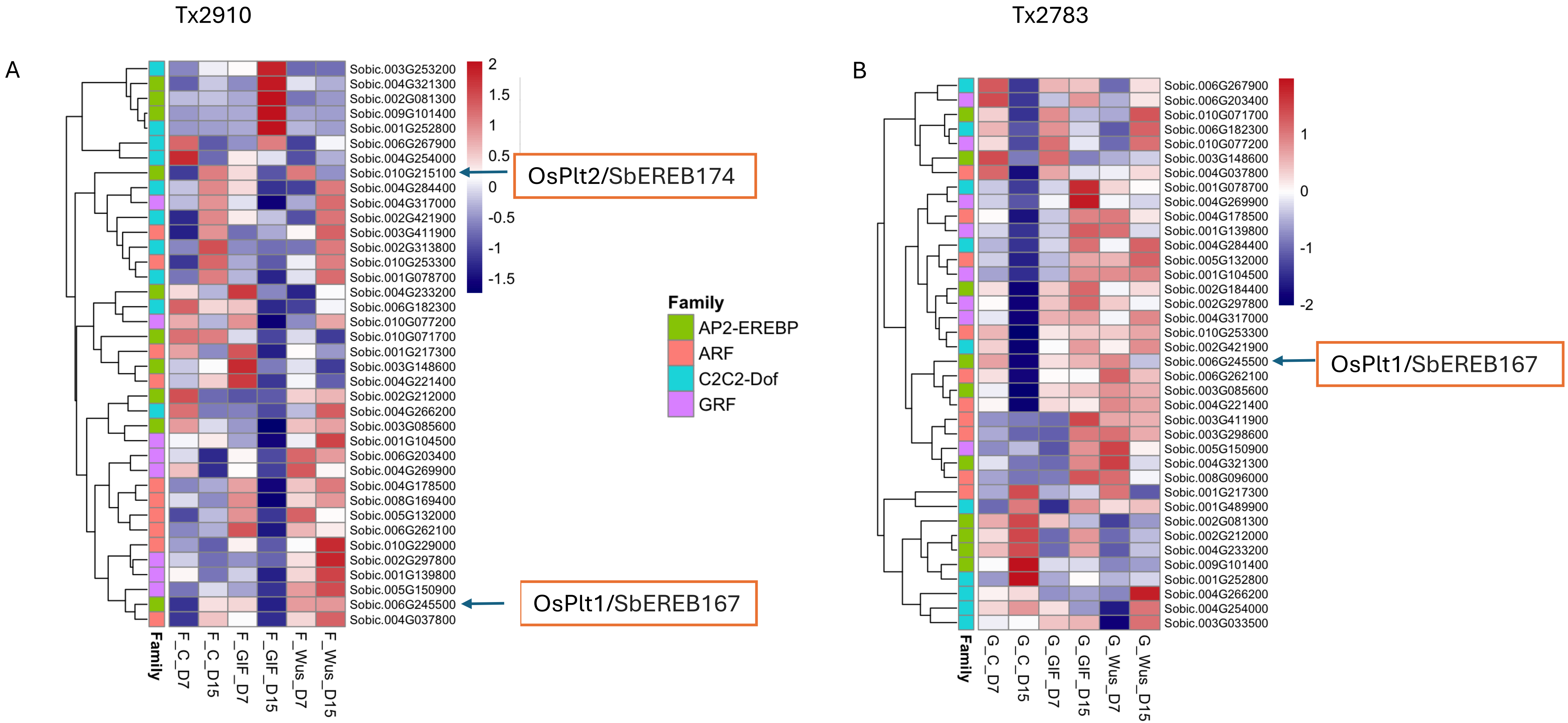


**Figure S11. Expression patterns of transcription factor (TF) families during somatic embryogenesis.** Mean Z-scaled expression of TF families across the RNA-seq samples of Tx2910 (A) and Tx2783 (B), with heatmaps of the top 10 members of selected families. *SbEREB174* (OsPLT2 ortholog) and *SbEREB167* (OsPLT1 ortholog) are labelled.

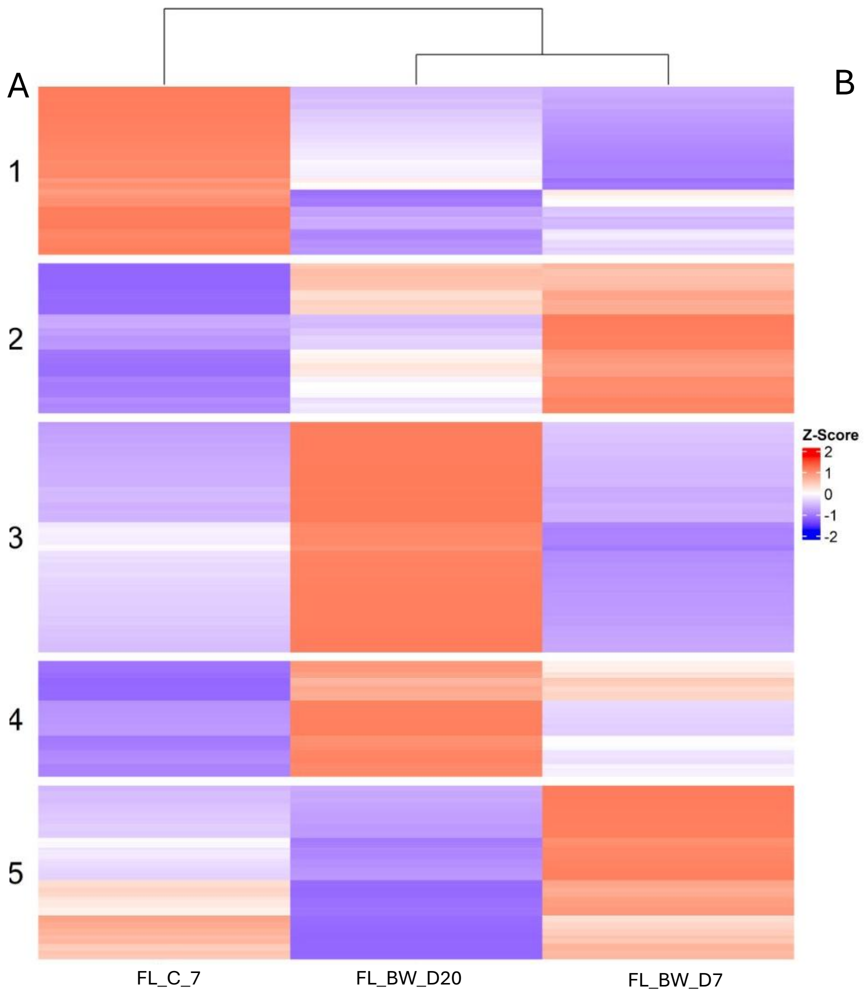


B

**GO biological Processes**

**Cluster 1**
Cutin biosynthesis
Maintenance of cell location
Phosphatydilcholine metabolic process
Amine biosynthesis

**Cluster 2**
Anatomical structure
Cell division/maturation
Hormone metabolic process
Phyto-steroids biosynthesis
Microtubules-based process

**Cluster 3**
Cell maturation
Xylem and phloem histogenesis
DNA replication
Specification of Symmetry
Regulation of phosphorylation

**Cluster 4**
Regulation of phosphorylation
Auxin mediate signaling
Spindle assembly
Pigment accumulation
Export across plasma membrane

**Cluster 5**
Cell surface receptor signaling
Jasmonic acid and ethylene systemic resistance
Flavonoids metabolic process
Aminoglycan/Amino sugar metabolic process

C

**Hub TFs**

**Cluster 1**
SbbZIP42, SbbZIP8,
SbMYB92

**Cluster 2**
SbGRAS19, SbGLK30,
SbOrphan23, SbABI28,
SbMYBR31

**Cluster 3**
SbABI19, SbABI33,
**SbGRF1, SbGRAS74,**
SbEREB22, SbEREB43,
SbGLK1, SbHSF12,
SbHSF9

**Cluster 4**
SbABI1, SbARF21,
SbbHLH52, SbGATA8,
SbJMJ11

**Cluster 5**
SbHB11, SbMYB60,
SbNAC10, SbWRKY89

**Figure S12. K-means transcriptional modules responding to ZmBBM + ZmWUS in forage sorghum (Tx2910) leaf explants.** Transcriptional modules identified by k-means clustering in forage sorghum (Tx2910) leaf explants. **A**, Heatmaps of DEGs (log2 fold change ≥ 2; Benjamini–Hochberg-adjusted P < 0.05) clustered into five clusters across the control at 7 DAT (FL_C_D7) and *ZmBBM* + *ZmWUS* at 7 and 20 DAT (FL_BW_D7, FL_BW_D20). **B**, Representative enriched GO biological-process terms for each cluster. **C**, Representative hub TFs for each cluster.

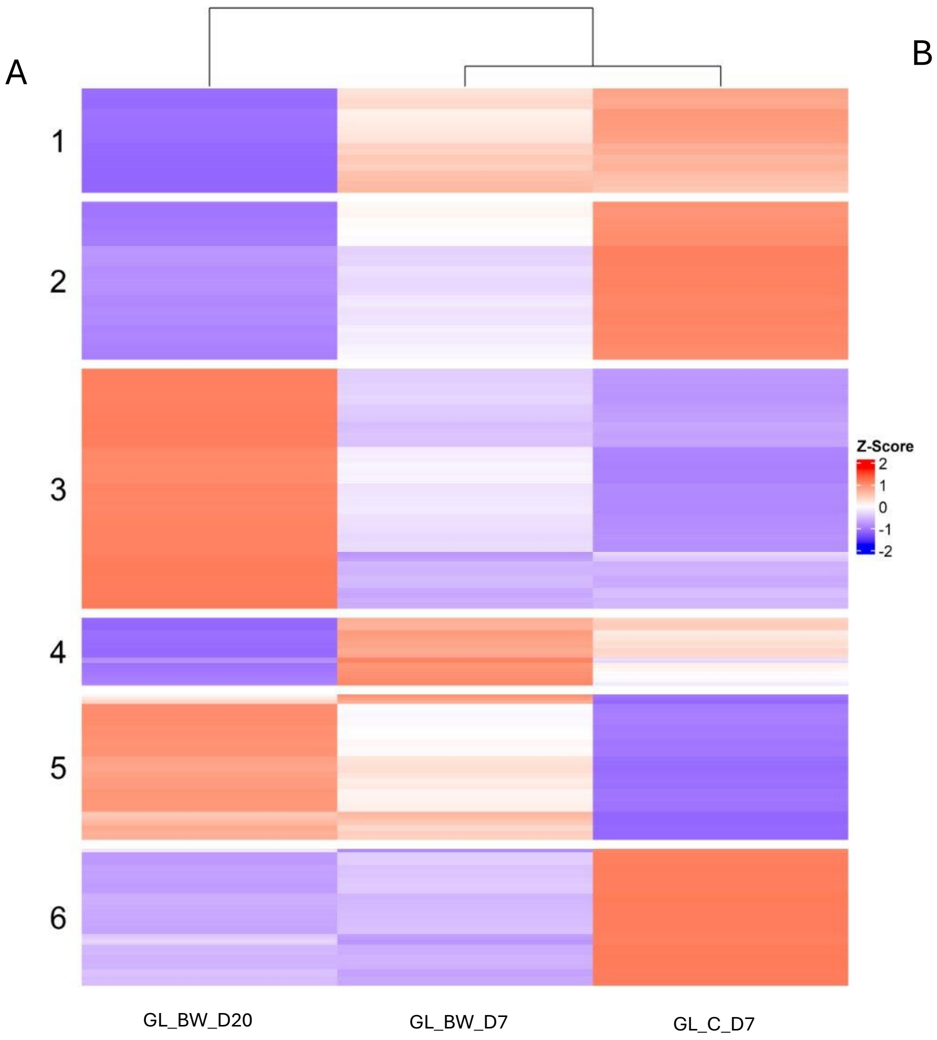


B **GO biological Processes**

**Cluster 1**
Flavonoid biosynthesis
Cutin biosynthesis
Phenylalanine metabolic process
Gibberellin metabolic process

**Cluster 2**
Anthocyanin containing compounds
Gibberellin metabolic process
Diterpenoid metabolic process
Cell wall assembly
Exocytosis

**Cluster 3**
Cell cycle phase transition
Sepal and Petal morphogenesis
Tyrosine kinase signaling

**Cluster 4**
Epidermis development
Carbohydrates biosynthesis
Metabolites and energy precursors
Photosynthesis regulation

**Cluster 5**
Cell cycle phase transition
Gibberellin metabolic process
IMP biosynthesis
Oligosaccharides

**Cluster 6**
Immune receptors
Response to peptide hormones
De-Etiolation
Trehalose metabolism
Starch catabolism

C **Hub TFs**

**Cluster 1**
SbBHLH103,
SbBHLH123, SbDOF23,
SbNAC33 ,

**Cluster 2**
SbEREBs,
SbHB46, SbNAC102, SbWRKY42,
SbZIM19, SBZIM5

**Cluster 3**
SbARF21, **SbBHLH1,** SbDOF17,
SbGRAS74, **SbGRF1, SbEREB174,**
**SbEREB165,**
**SbGRF6, SbHB44,** SbHB69,
SbNAC95, SbEREB153, **SbHB44**

**Cluster 4**
SbbZIP16, SbHB60,
SbMYB48

**Cluster 5**
SbEREB45, SbARF27,
SbYAB2, SbMYBR71,
SbHB32 (WUS related)

**Cluster 6**
SbZIM14, SbZIM15,
SbZIM9, SbWRKY29,
SbMYB7, SbHSF4

**Figure S13. K-means transcriptional modules responding to ZmBBM + ZmWUS in grain sorghum (Tx2783) leaf explants.** Transcriptional modules identified by k-means clustering in grain sorghum (Tx2783) leaf explants. **A**, Heatmaps of DEGs (log2 fold change ≥ 2; Benjamini–Hochberg-adjusted $P < 0.05$) clustered into six clusters across the control at 7 DAT (GL_C_D7) and *ZmBBM* + *ZmWUS* at 7 and 20 DAT (GL_BW_D7, GL_BW_D20). **B**, Representative enriched GO biological-process terms for each cluster. **C**, Representative hub TFs for each cluster.

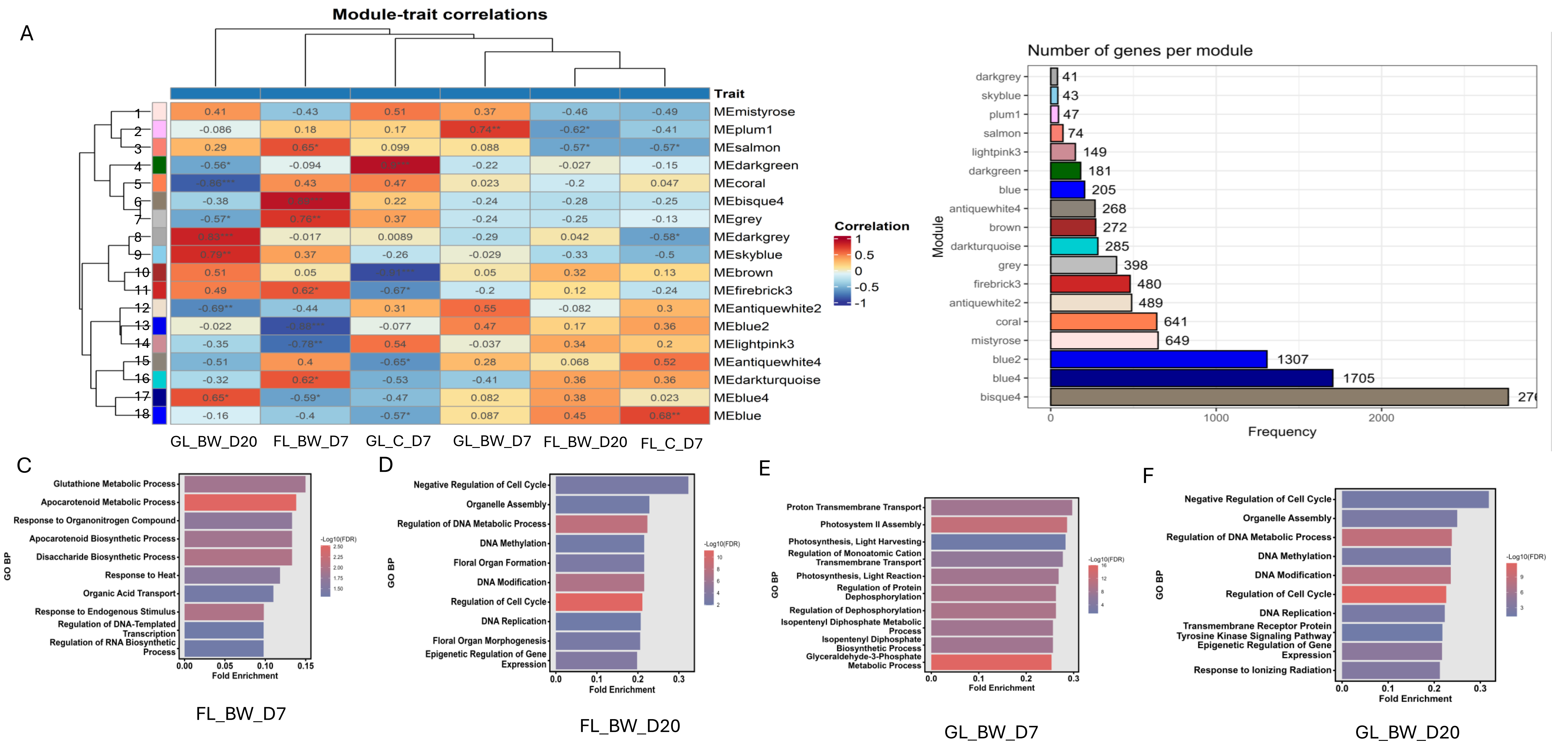


**Figure S14. Weighted gene co-expression network analysis (WGCNA) of leaf explants. A**, Module–trait correlation heatmap for the 18 co-expression modules and the six leaf samples (FL_C_D7, FL_BW_D7, FL_BW_D20, GL_C_D7, GL_BW_D7, GL_BW_D20); the modules fall into three categories of sample-associated expression programs. **B**, Number of genes in each module. **C–F**, GO biological-process enrichment of the modules associated with FL_BW_D7 (C), FL_BW_D20 (D), GL_BW_D7 (E) and GL_BW_D20 (F).

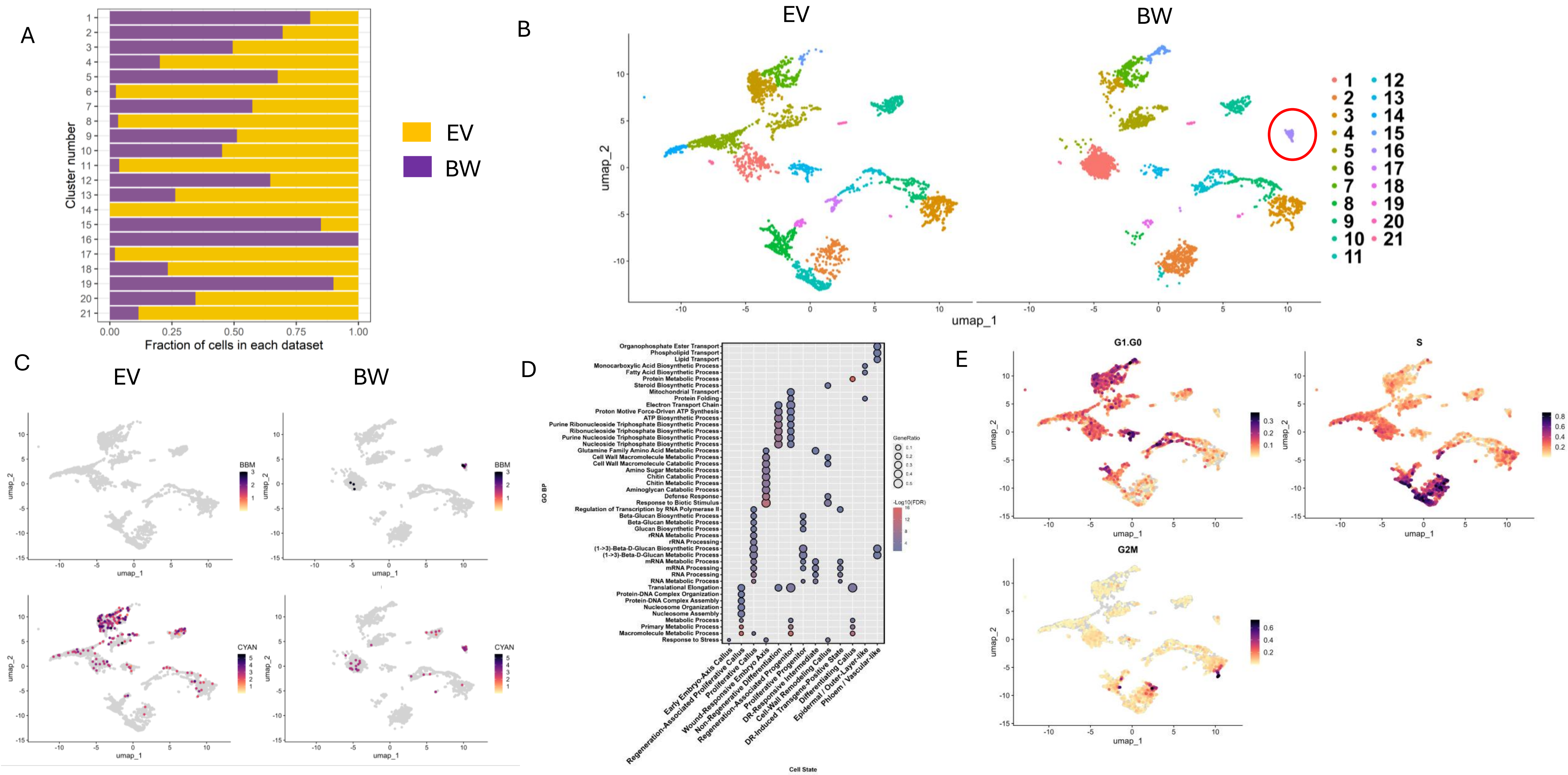


**Figure S15. Single-nucleus RNA-seq of EV and ZmBBM + ZmWUS (BW) tissue. A**, Fraction of nuclei from EV and BW samples in each of the 21 clusters. **B**, UMAP of the integrated dataset split by sample, coloured by cluster; the red circle marks a BBM + WUS -specific cluster. **C**, Feature plots of transgene expression (*ZmBBM* and *AmCyan*) split by sample; darker colour indicates higher expression. **D**, GO biological-process terms enriched among cluster marker genes (FDR < 0.05). **E**, Feature plots of cell-cycle phase scores (G1/G0, S, G2/M); darker colour indicates higher scores.

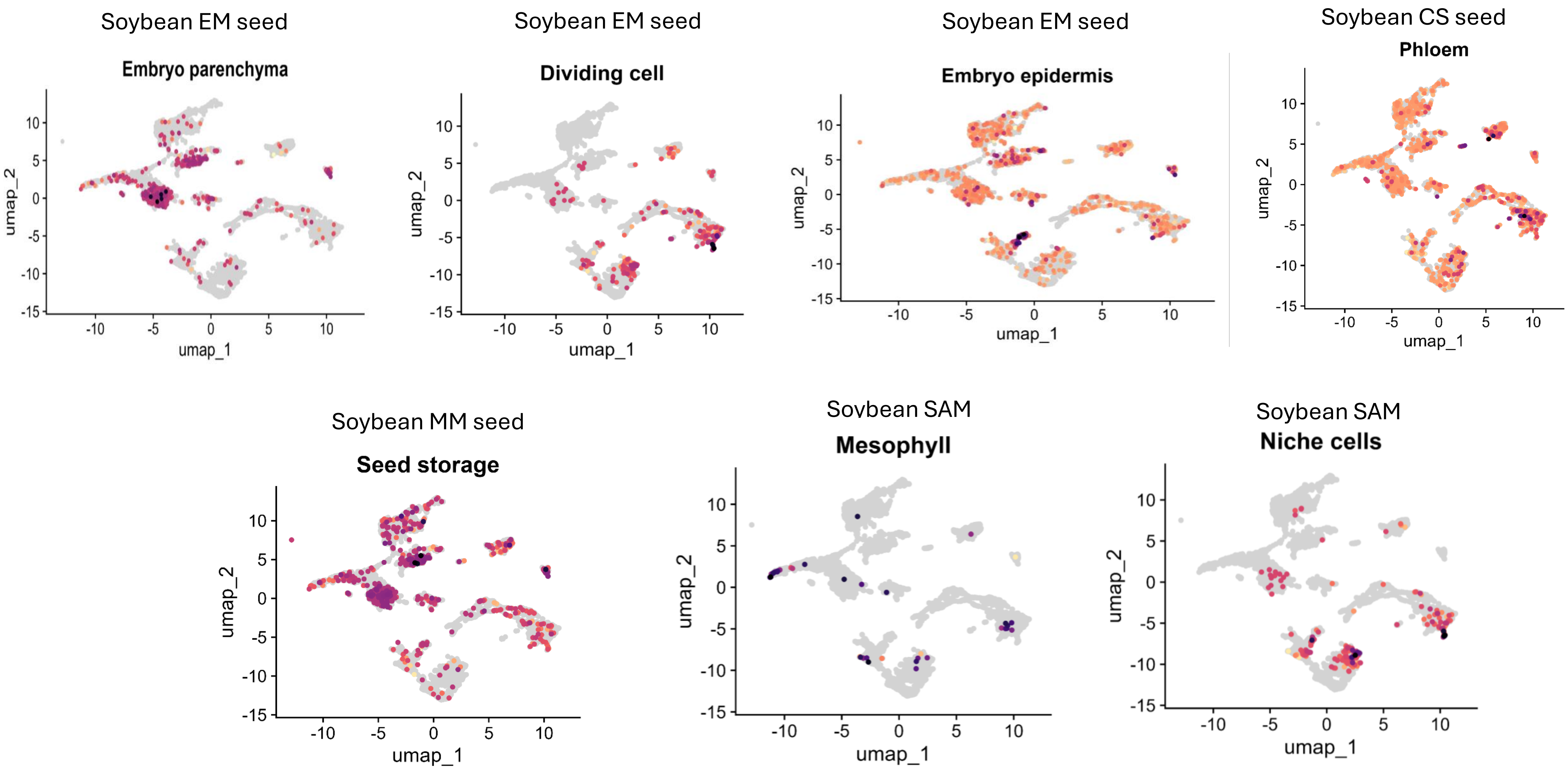


**Figure S16. Expression of soybean cell-type marker orthologs in the sorghum snRNA-seq dataset.** UMAP feature plots showing expression of marker genes for cell types defined in published soybean seed and shoot apical meristem (SAM) snRNA-seq datasets: early-maturity (EM), cotyledonary-stage (CS) and middle-maturity (MM) seeds, and SAM. Source datasets:doi:10.1016/j.cell.2024.10.050 and bioRxiv, doi:10.1101/2024.07.08.602332. Marker genes are listed in Supplementary Data.

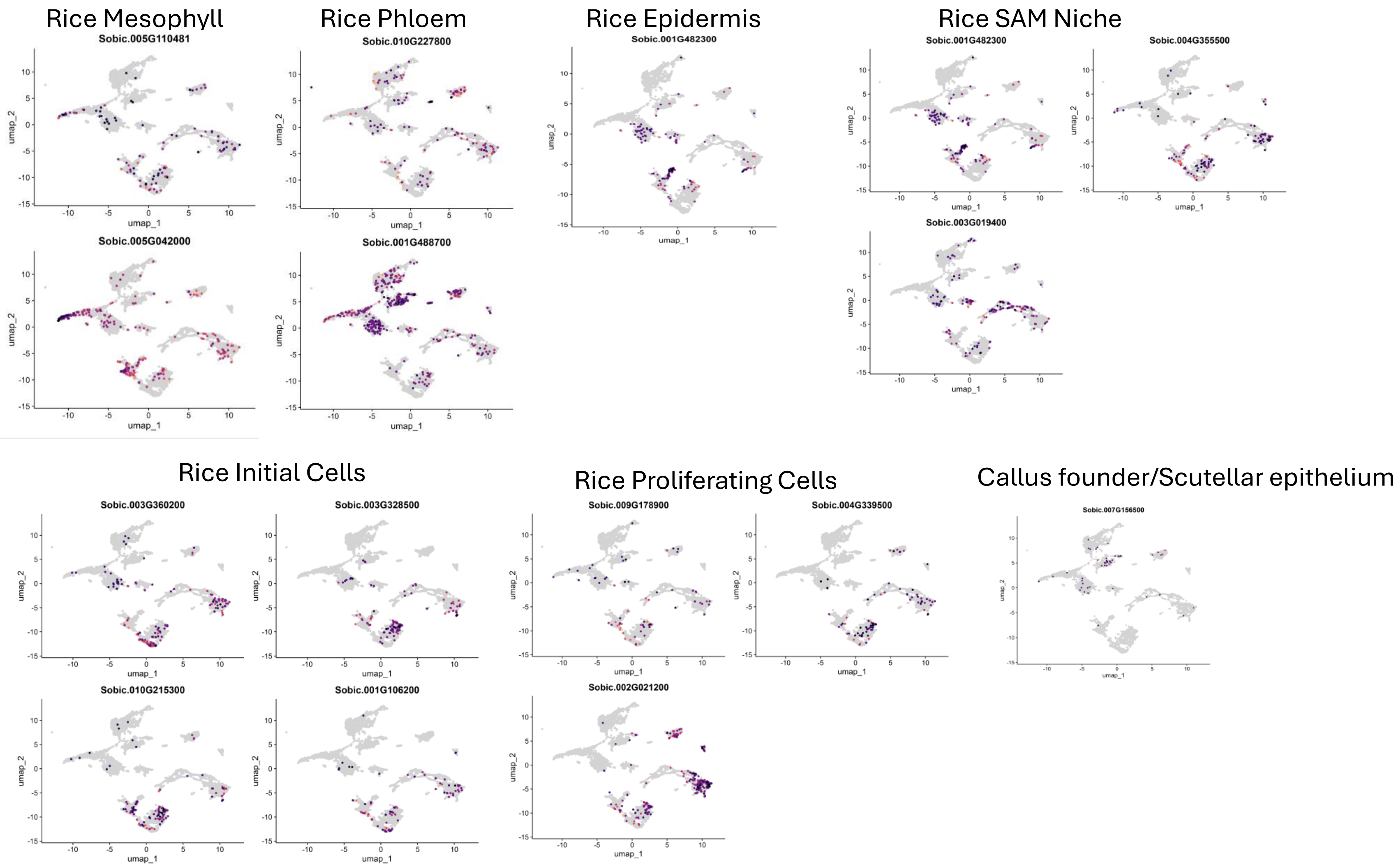


**Figure S17. Expression of rice cell-type marker orthologs in the sorghum snRNA-seq dataset.** UMAP feature plots showing expression of marker genes for rice cell types: mesophyll, phloem, epidermis, callus founder cell/scutellar epithelium, initial cells, proliferating cells and SAM niche. Source dataset: doi:10.1186/s12870-026-08596-6. Marker genes are listed in Supplementary Data.

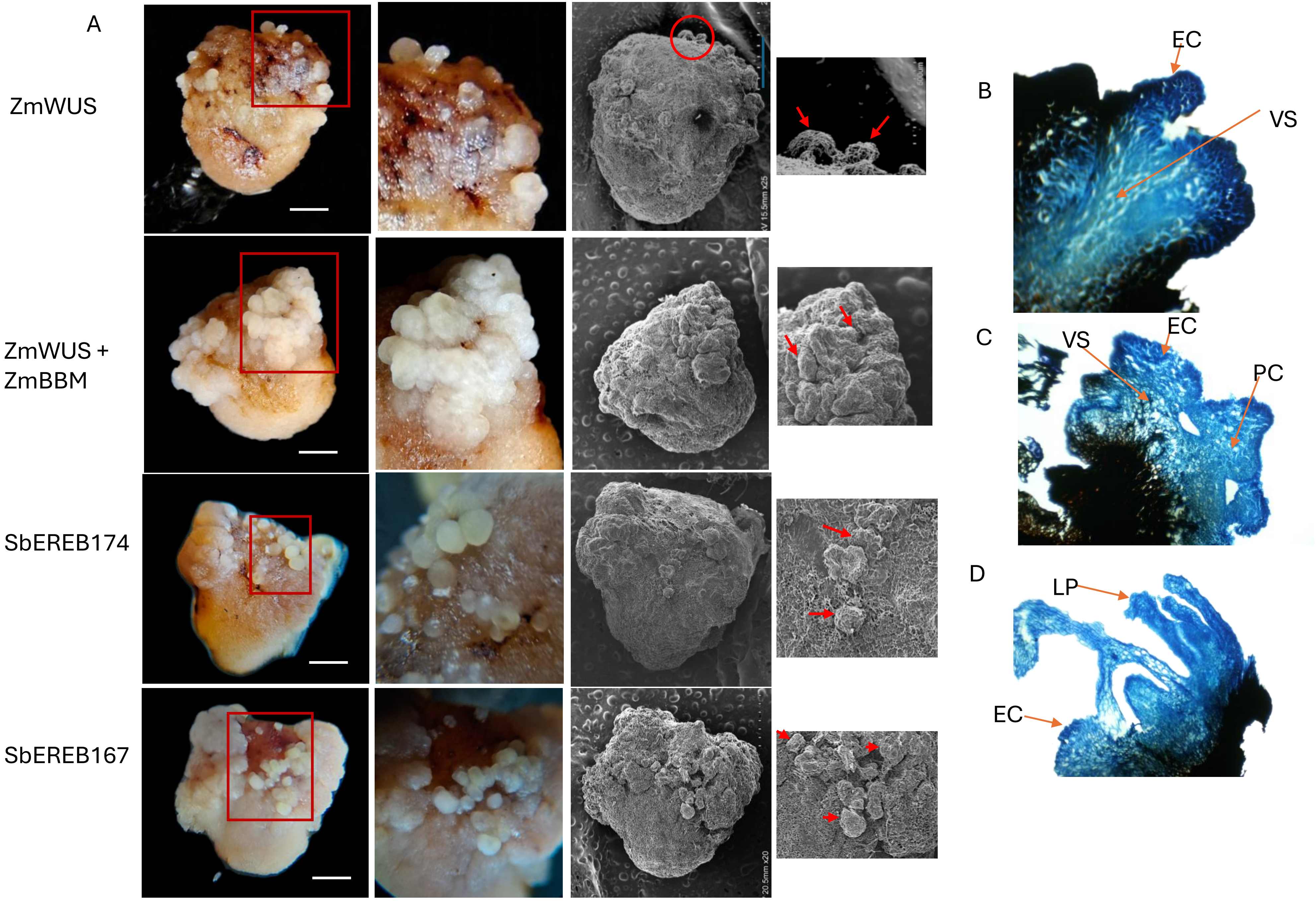


**Figure S18. Morphology and histology of somatic embryogenesis induced by DRs.**

**A**, Tx2783 IEs expressing *ZmWUS*, *ZmWUS* + *ZmBBM*, *SbEREB174* or *SbEREB167* at 7 DAT, with scanning electron microscopy (SEM) images of the corresponding embryogenic structures. Scale bar, 1 mm. **B–D**, Histological sections of *SbEREB174*-overexpressing calli at early (B), intermediate (C) and late (D) embryogenic stages.

EC, embryogenic cells; PC, parenchymatous callus cells; VS, vasculature; LP, leaf primordia.

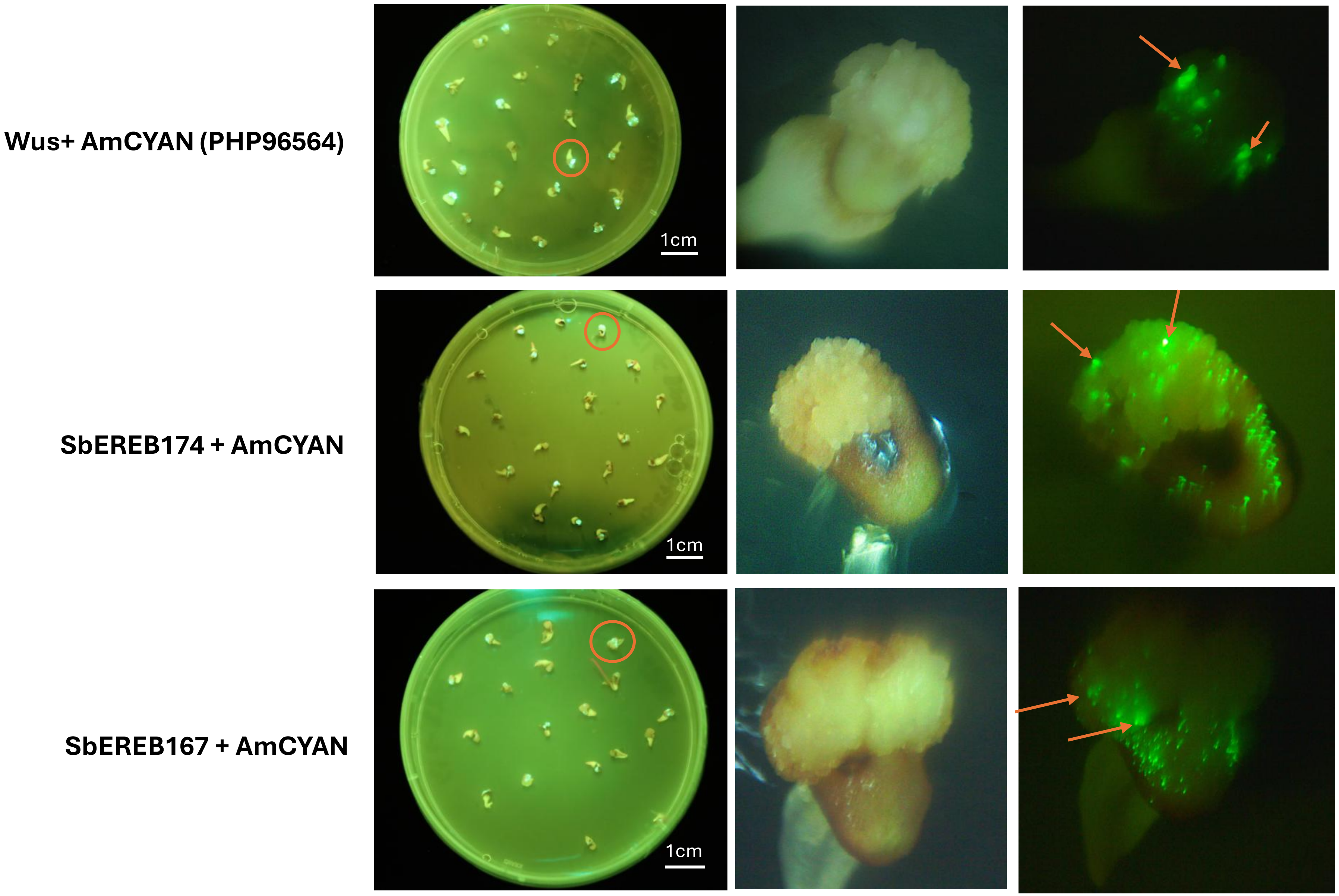


**Figure S19. Early detection of embryogenic foci with the AmCYAN marker.** Tx2783 IEs expressing *ZmWUS* + AmCYAN (PHP96564), *SbEREB174* + AmCYAN or *SbEREB167* + AmCYAN at 12 DAT. AmCYAN fluorescence marks transgenic callus and enables early detection of embryo formation; arrows indicate foci of morphogenic structure development.

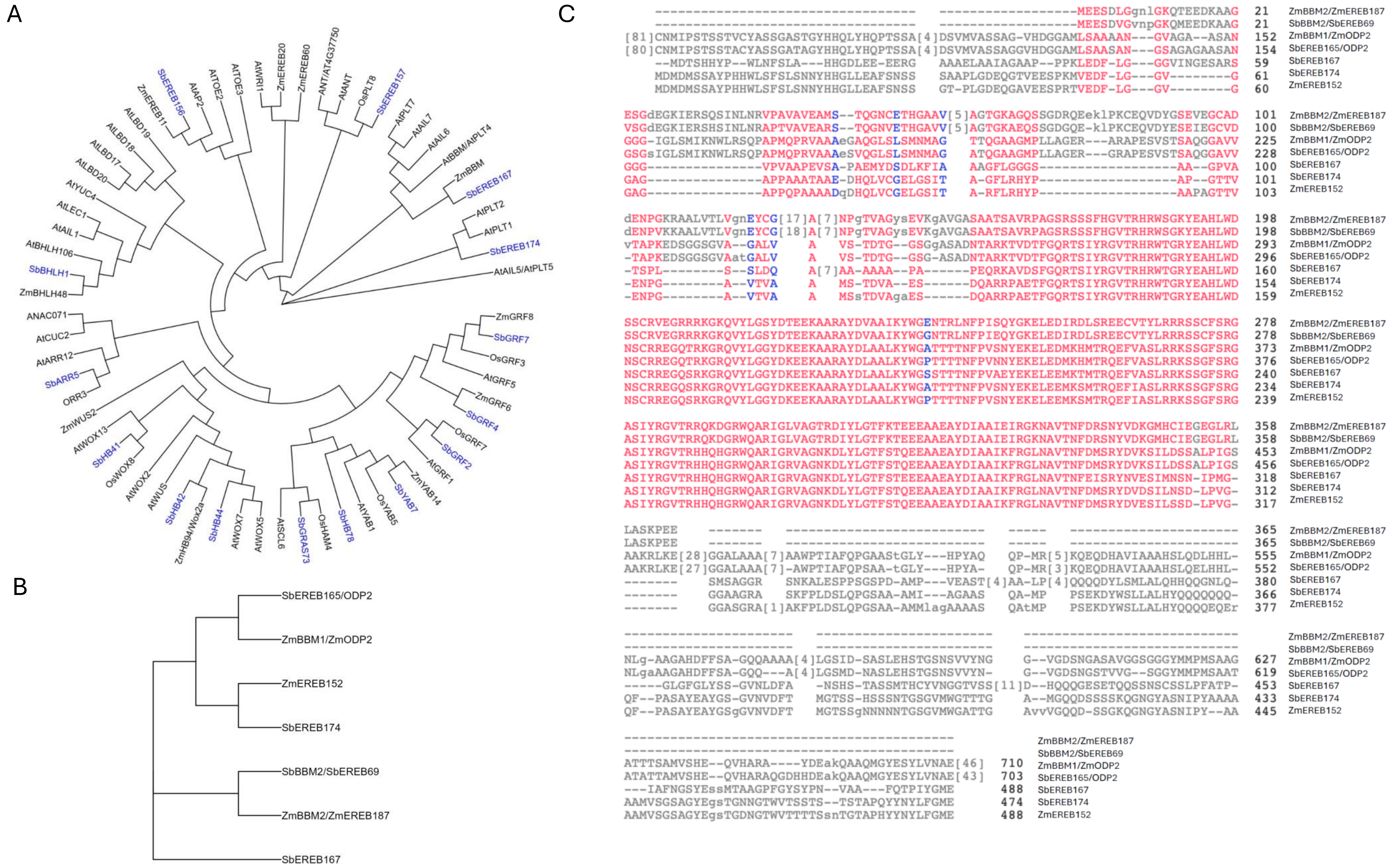


**Figure S20. Phylogeny and sequence alignment of the 15 candidate regulators. A**, Maximum-likelihood tree of the 15 sorghum candidates (blue) with genes associated with pluripotency in Arabidopsis, rice and maize. **B**, Tree of AP2/EREBP-family genes and related developmental regulators. **C**, Sequences of the sorghum AP2/EREBP candidates and related regulators are shown relative to the reference sequence (ZmBBM2/ZmEREB187). Protein sequences were aligned with MUSCLE and trees were inferred by maximum likelihood in R.

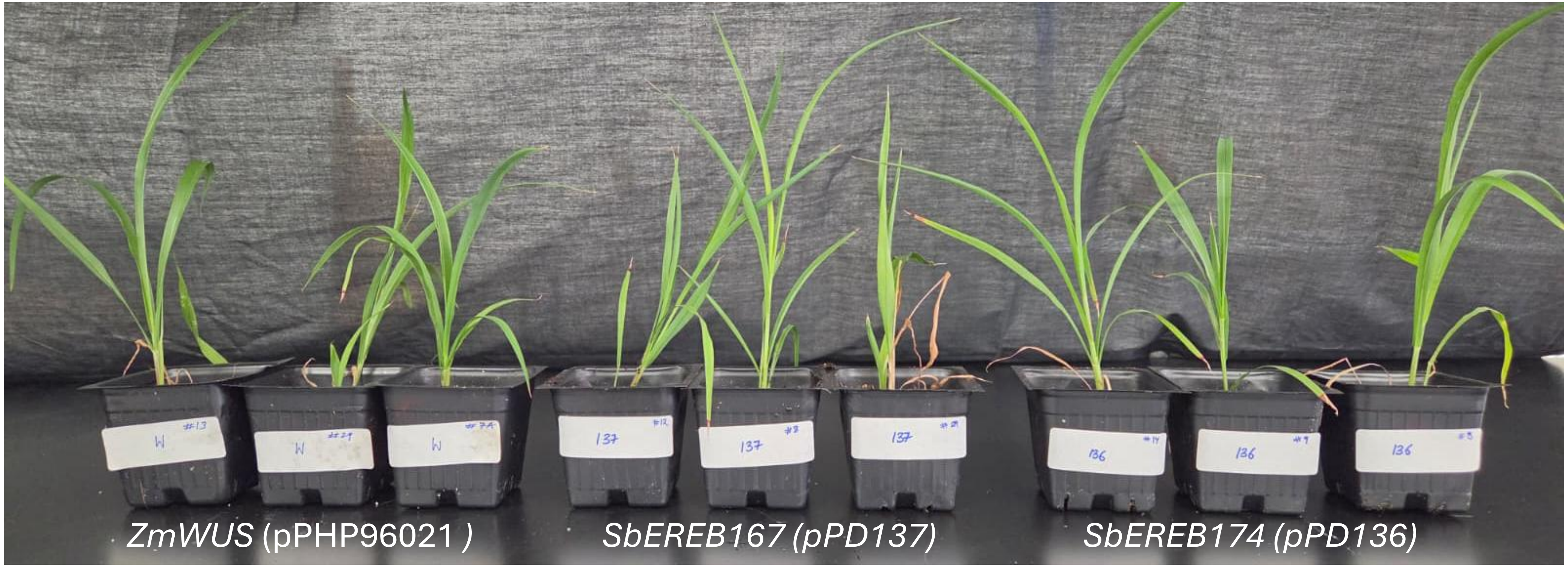


**Figure S21: Morphology of representative $T_0$ transgenic plants after transfer to soil.** Tx2783 plants regenerated from immature embryos expressing *ZmWUS* (positive control; label "W"), *SbEREB167* (pPD137; label "137") or *SbEREB174* (pPD136; label "136"). Three independent events per construct are shown. Plants were photographed after hardening in soil (2 weeks after transfer). All plants show normal leaf and shoot morphology without or deformed shoots.
